# Mitochondrial Double-Stranded RNA Triggers Ferroptosis via PKR–ISR Signaling in Renal Ischemia–Reperfusion Injury

**DOI:** 10.64898/2026.01.19.700463

**Authors:** Richie Justin, Maria Sophia Criste, Soyoung Park, Fedho Kusuma, Nara Lee, Yerin Yang, Kim Anh Nguyen, Samel Park, Jeongan Kim, Hyo-Wook Gil, Jaeseok Han

## Abstract

Renal ischemia–reperfusion (IR) injury remains a leading cause of acute kidney injury, although the precise molecular pathways responsible for tissue injury are not fully understood. In this study, we demonstrate that IR initiates ferroptosis through activation of the PKR-dependent integrated stress response (ISR), which is driven by the accumulation of mitochondrial double-stranded RNA (dsRNA). Our mechanistic investigations reveal that IR perturbs mitochondrial homeostasis, leading to the accumulation of mitochondrial dsRNA and its subsequent release into the cytosol, thereby activating the dsRNA-dependent kinase PKR. Activation of PKR results in eIF2α phosphorylation and maladaptive ISR signaling, which facilitates ferroptosis and impairs renal function. Notably, genetic ablation of PKR or pharmacological intervention to inhibit ISR significantly reduced ferroptosis and preserved renal function. Collectively, these results identify the mitochondrial dsRNA–PKR–ISR axis as a critical mediator of ferroptotic renal injury and highlight its potential as a therapeutic target in ischemic acute kidney injury.

## Main

Ischemia-reperfusion (IR) injury in the kidney is a primary cause of acute kidney injury (AKI), frequently occurring in clinical scenarios such as renal transplantation, sepsis, or major surgeries^1,2^. This syndrome is characterized by a rapid decline of renal function, which leads to compromised clearance of metabolic waste and disruption of fluid and electrolyte balance^2^. Epidemiological data indicate that the incidence of AKI ranges from 5–20% among all hospitalized patients, increasing to 20–50% for those admitted to intensive care units, where IR injury constitutes the predominant etiology^3–5^. AKI is frequently reported to progress to chronic kidney disease and is recognized as a major risk factor for the development of end-stage renal disease^6^. The underlying mechanisms of renal IR injury are multifaceted, with significant contributions from oxidative stress, inflammatory processes, and regulated cell death^7^. While restoration of blood flow through reperfusion is necessary for tissue survival, it paradoxically intensifies tissue injury by promoting a surge in reactive oxygen species (ROS) production, mitochondrial dysfunction, and metabolic derangements^8,9^. Despite extensive research efforts, the specific signaling pathways linking IR to subsequent renal injury have yet to be fully elucidated.

Mitochondria are double-membrane organelles responsible for ATP production through oxidative phosphorylation^10^. Additionally, they regulate redox/ROS homeostasis, buffer Ca²⁺, and generate biosynthetic precursors^11^. Mitochondria also maintain biosynthetic flux and cell-fate determination through the processes of fission–fusion dynamics, mitophagy, and biogenesis^10,12^. These functions position mitochondria as key integrators of metabolic and stress signals in renal tubular cells, while, under injurious conditions, making them the most susceptible organelles that play a critical role in determining the severity of IR injury^8,13,14^. During ischemia, loss of mitochondrial membrane potential and ATP depletion disrupt cellular homeostasis^13,15^. Upon reperfusion, the abrupt restoration of oxygen triggers excessive ROS production, impairs mitochondrial complexes, induces mitochondrial permeability transition pore (mPTP) opening, and causes cytochrome c release, together promoting apoptotic and necrotic cell death^16–18^.

When exposed to these stresses, injured mitochondria trigger adaptive mechanisms that include alterations in mitochondrial dynamics, initiation of mitophagy, and transmission of retrograde signals to the nucleus^19–21^. Although an imbalance in mitochondrial dynamics and malfunctioning mitophagy have been identified as pivotal early factors that intensify renal tubular damage following reperfusion^18^, the subsequent mitochondrial signaling events during IR injury remain incompletely characterized. When mitochondrial function is disrupted, cells initiate mitochondrial retrograde signaling, allowing stressed mitochondria to relay their status to the cytosol and nucleus to coordinate either adaptive or maladaptive responses^22,23^. These signaling pathways engage cytosolic and nuclear sensors, prompting a reprogramming of transcription, translation, and metabolic activity aimed at reestablishing cellular homeostasis^22^. Among these pathways, the integrated stress response (ISR) has been especially well characterized, in which stress-responsive kinases such as PERK, GCN2, HRI, and PKR phosphorylate eIF2α to temporarily inhibit global protein synthesis while promoting ATF4-dependent transcriptional programs involved in the regulation of antioxidant defense, metabolism, and cell survival^24–26^. Although ISR has been implicated in a variety of pathological conditions, including neurodegenerative and metabolic diseases^25,27^, its specific role in the IR process, particularly in renal IR, has not yet been fully elucidated.

Recent studies have identified ferroptosis, a regulated form of cell death characterized by iron-dependent lipid peroxidation, as a major factor in IR-associated diseases. It has been widely investigated in organs such as the brain, heart, liver, and kidney^28–31^. Elevated mitochondrial ROS and impaired antioxidant systems, especially those involving glutathione and GPX4, promote lipid peroxide accumulation, thereby increasing cellular susceptibility to ferroptosis during IR injury^32–34^. Mitochondrial dysfunction has been demonstrated to exacerbate damage in renal IR injury, as impairment of the antioxidant defense system fails to control excessive ROS generation, which is believed to drive the lipid peroxidation process leading to ferroptosis^28,35^. The relationship between mitochondrial dysfunction, ISR activation, and ferroptosis forms a crucial axis influencing renal cell fate under IR conditions. In this context, we introduce a comprehensive molecular pathway linking mitochondrial stress, ISR activation, and ferroptosis in renal IR injury. Elucidation of this interconnected network may reveal innovative therapeutic options for reducing renal injury in IR-associated AKI.

## Results

### IR causes activation of ISR in kidney

IR impairs mitochondrial function^17,36^ and may initiate retrograde signaling pathways that activate the ISR^37^. To clarify this association, we examined a published transcriptomic dataset (GSE267650) generated from mouse kidneys exposed to 20 minutes of ischemia followed by 16 hours of reperfusion^38^. Many of the differentially expressed genes identified in this analysis were linked to mitochondrial function disturbances (Extended Fig. 1a). Gene set enrichment analysis (GSEA) indicated substantial downregulation of oxidative phosphorylation and tricarboxylic acid (TCA) cycle pathways (Fig. 1a), while gene ontology (GO) analysis corroborated disturbances in mitochondrial biological processes (Extended Fig. 1b). Notably, mitochondrial dysfunction became more pronounced with increased reperfusion duration, but was not detected following ischemia alone (Extended Fig. 1c). To extend our observations, we analyzed transcriptomic data from human kidneys derived from a public dataset (GSE126805), including samples from patients undergoing kidney transplantation as the underlying cause of renal ischemia–reperfusion (IR) injury^39^. Comparative analysis of shared differentially expressed genes between mouse and human datasets revealed both enrichment and elevated expression of ISR-related transcripts, and this transcriptional adaptation was further validated by pathway-level activation observed in the violin plot analysis (Fig. 1b). Building upon recent findings, including our own, which demonstrate that mitochondrial dysfunction can activate the ISR^37,40,41^, we observed a marked induction of canonical ISR target genes (*Ddit3, Hspa1b, Atf5, Hspb1, Ppp1r15a, Atf3, Slc2a1, Hspd1*)^42,43^ following renal IR (Fig. 1c), supporting a mechanistic link between mitochondrial disruption and stress-responsive signaling. Correlation network analysis identified strong positive correlations among ISR genes, indicating a coordinated activation program in response to IR-induced mitochondrial stress (Extended Fig. 1d). To further validate these findings, we modeled IR injury in human renal proximal tubule epithelial cells (HK-2) using hypoxia–reoxygenation (H/R)^44^ and quantified mRNA levels of genes implicated in the ISR pathway. In agreement with the transcriptomic data, H/R exposure led to a significant upregulation of these genes (Fig. 1d). This increase in transcript levels was accompanied by phosphorylation of eIF2α, a key event in ISR activation (Fig. 1e).

**Fig. 1:**
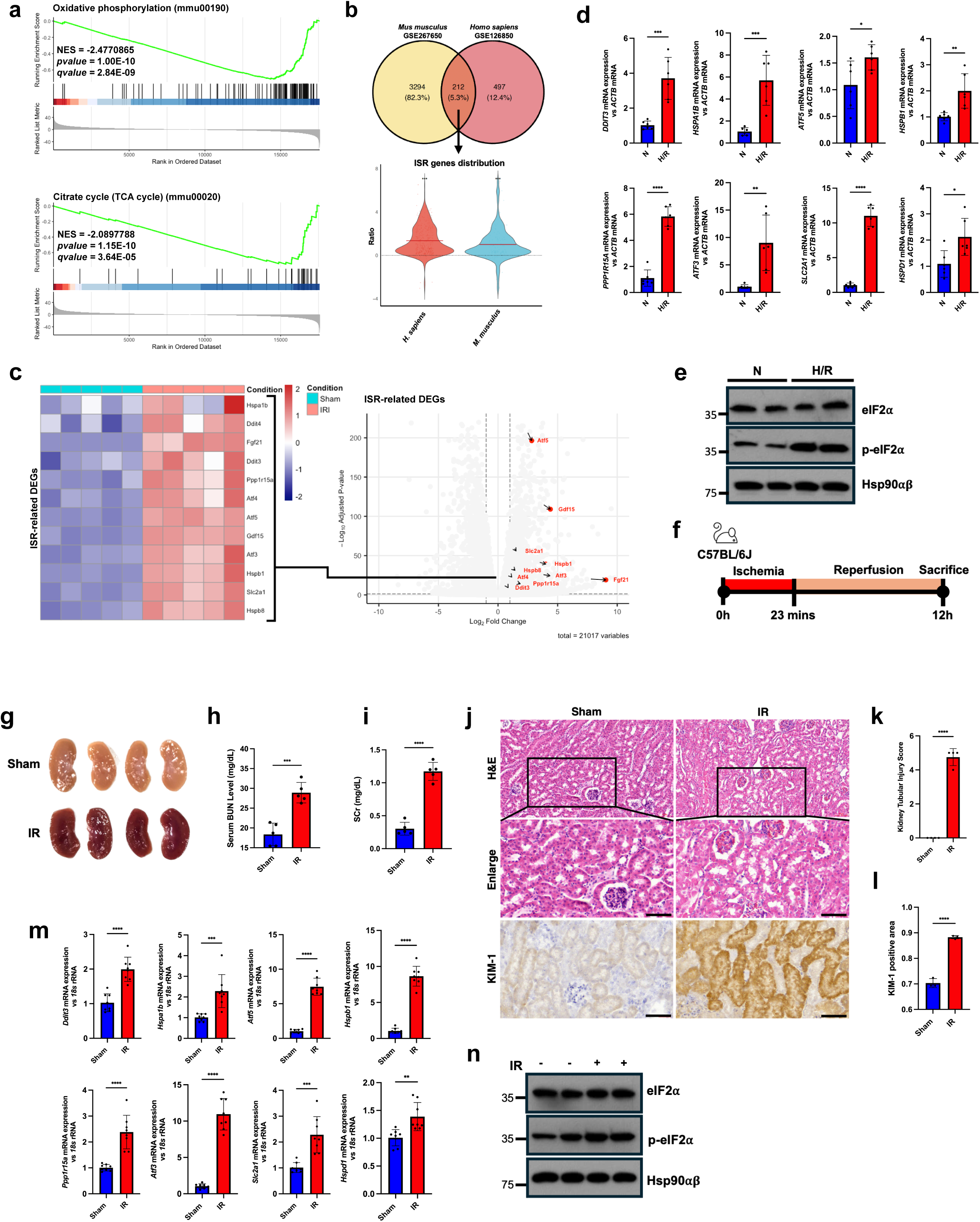
Ischemic-reperfusion (IR) injury induces activation of the integrated stress response (ISR) in the kidney. **a,** Gene set enrichment analysis (GSEA) using the RNA-Seq dataset (GSE267650) from mice subjected to 20 mins ischemia followed by 16 hours reperfusion was performed focusing on reactome signaling pathways: oxidative phosphorylation and Citrate cycle (TCA cycle) in the context of kidney injury induced by IR. The enrichment analysis revealed significant downregulation of these pathways in IR-treated mice compared to sham-treated controls, indicating a mitochondrial dysfunction signature typical of renal IR injury. **b,** Venn diagram (top) showing the overlap between differentially expressed genes from mouse kidneys subjected to 20 min ischemia followed by 16 h reperfusion (GSE267650) and paired human kidney biopsy transcriptomes collected before and after transplantation reperfusion (GSE126805). The shared gene set was further filtered for ISR-related transcripts and used to generate the violin plot (bottom). In the violin plot, each dot represents the log_10_-normalized expression value of an individual ISR-associated gene; violin width reflects expression density and center lines denote the mean. **c,** The heatmap visualizes the increased expression of ISR-associated differentially expressed genes between sham-treated and IR-induced kidney injury in mouse transcriptomic data. The colored scale bar at the upper right of the heatmap represents relative expression values, where −2 and 2 denote under- and over-abundance, respectively. **d,** RT-qPCR was performed to analyze ISR-related mRNA expression following 6 hours hypoxia and 1 hour reperfusion in HK-2 cells. *N* = *6.* **e,** After exposing HK-2 cells to 6 hours hypoxia and 1 hour reperfusion, protein levels of eIF2⍺ and p-eIF2⍺ were assessed by immunoblotting. Hsp90 was used as the protein loading control. **f,** Schematic representation of the experimental protocol for inducing bilateral IR kidney injury in mice. **g,** Macroscopic morphological comparison of sham- and IR-induced kidneys. Images were acquired immediately post-mortem after whole body perfusion via the heart. **h-i,** Assessment of serum BUN and creatinine levels between mice subjected to sham and kidney IR injury procedures. *N = 5*. **j-l,** Representative H&E- and KIM-1-immunohistochemistry-stained kidney sections. Tubular injury scores were determined by quantifying the percentage of tubules exhibiting kidney injury-associated morphology as previously described, including regions of dilatation, desquamation, vacuolization, necrosis, atrophy, casts, interstitial inflammatory cell infiltration, and edema. The KIM-1-positive area was measured by normalizing chromogenic intensity to hematoxylin. *N = 4*. Scale bar = 60 µm. **m,** RT-qPCR analysis of ISR-related mRNA expression in kidneys from sham- and IR-treated mice. *N* = *8.* **n,** Immunoblot analysis demonstrating the expression of eIF2⍺ and p-eIF2⍺ in mouse kidneys following IR procedure. Statistical analysis was conducted using a two-tailed unpaired t-test. *\*P < 0.05, **P < 0.01, ***P < 0.001, ****P < 0.0001*.

To further validate the bioinformatic and *in vitro* findings *in vivo*, C57BL/6J mice underwent 23 minutes of bilateral renal pedicle clamping followed by 12 hours of reperfusion (Fig. 1f), as described previously^45^. Gross examination of kidneys retrieved after reperfusion demonstrated pronounced tissue injury in the IR group (Fig. 1g). Renal function was significantly compromised, with serum blood urea nitrogen (BUN) and creatinine (SCr) levels markedly increased^46^ (Fig. 1h,i). Consistent with these biochemical changes, hematoxylin and eosin (H&E)-stained histology revealed substantial disruption of renal structure in the IR group (Fig. 1j,k), assessed as previously documented^45^. Morphological evaluation using aquaporin-1 (AQP-1) staining, a marker for proximal tubular cells^47^, showed substantial injury of proximal tubular cells (Extended Fig. 1e). Immunohistochemical analysis confirmed tubular damage through kidney injury molecule-1 (KIM-1), a well-established biomarker of renal pathology^48^, quantified via DAB chromogenic intensity (Fig. 1j,l). Correspondingly, both ISR-related gene expression (Fig. 1m) and eIF2α phosphorylation (Fig. 1n) were substantially elevated after IR, signifying activation of the ISR pathway in response to IR-induced cellular stress. Collectively, findings from cross-species, *in vitro*, and *in vivo* studies consistently demonstrate that renal IR injury leads to mitochondrial stress and subsequent activation of the ISR pathway.

### Inhibition of ISR preserves kidney from renal damage caused by IR

After observing robust activation of the ISR following renal IR injury, we proceeded to examine the functional role of ISR signaling in kidney injury using pharmacological modulators. To suppress ISR activity, we utilized ISRIB and 2BAct, which are selective eIF2B activators that restore protein translation independently of eIF2α phosphorylation^49,50^. In H/R-treated HK-2 cells, both compounds markedly suppressed the expression of ISR-related genes (Fig. 2a, Extended Fig. 2a). Furthermore, CCK-8 assays confirmed that ISR inhibition was associated with improved cell viability under H/R conditions, supporting a deleterious role of excessive ISR activation in cellular injury during H/R (Fig. 2b). Notably, treatment with 2BAct preserved eIF2α phosphorylation levels, which aligns with its mechanism of bypassing translational inhibition from eIF2α phosphorylation by directly enhancing eIF2B activity^49,50^ (Extended Fig. 2b).

**Fig. 2:**
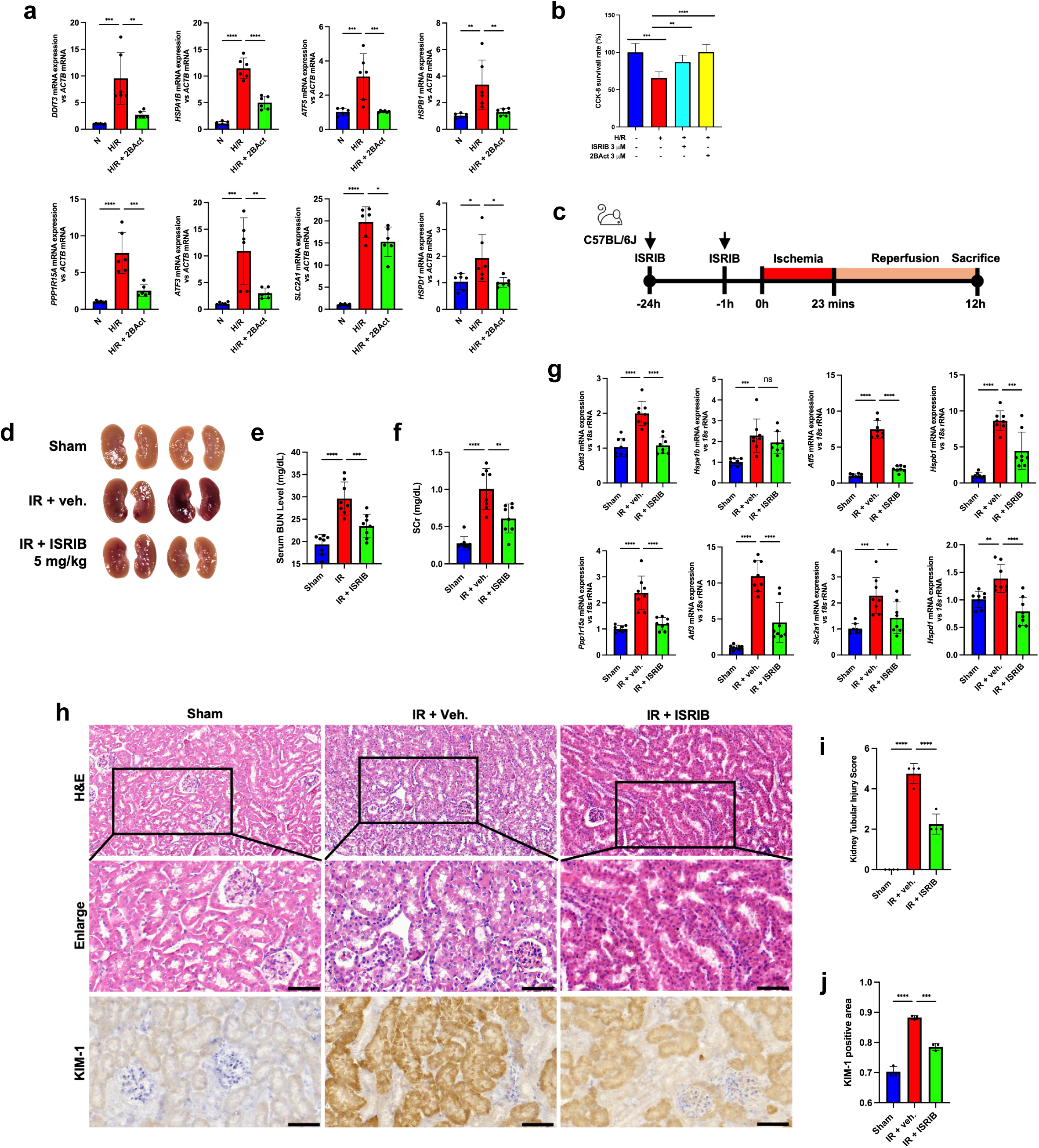
Inhibition of ISR preserves the kidney from renal damage caused by IR. **a,** Gene expression profiling to assess ISR-related mRNA levels after 6 hours of hypoxia and 1 hour of reperfusion, with or without 1 hour of pretreatment and co-treatment during reperfusion with 2BAct to inhibit ISR activation. *N = 6.* **b,** CCK-8 assay for cell viability following H/R exposure in HK-2 cells after ISR inhibition using ISRIB and 2BAct. *N =* 6. **c,** Schematic representation of the experimental design for intraperitoneal ISRIB administration in the renal IR-induced mouse model. **d,** Representative images showing macroscopic morphology in sham, IR, and ISRIB-treated mouse kidney. **e-f,** Quantification of serum BUN and SCr levels in mice subjected to sham or kidney IR procedures. *N = 8.* **g,** RT-qPCR quantification of ISR-related mRNA expression comparing sham, IR, and ISRIB-treated mice. *N* = 8. **h-j,** Histological evaluation using H&E staining and KIM-1 immunohistochemistry on kidney sections. The tubular injury score is determined by assessing the percentage of tubules exhibiting kidney injury-associated changes as previously described. The KIM-1 positive area is quantified by normalizing chromogenic signal intensity to hematoxylin. *N =* 4. Scale bar = 60 µm. Two-tailed unpaired t-test was used for data analysis (**b**), and one-way ANOVA for multiple comparisons, unless otherwise specified. *\*P < 0.05, **P < 0.01, ***P < 0.001, ****P < 0.0001*.

Based on these *in vitro* results, we then explored the impact of ISR signaling *in vivo* by administering ISRIB intraperitoneally to mice 24 hours and 1 hour prior to IR surgery (Fig. 2c). Following ISRIB administration, gross examination revealed reduced structural damage in kidneys (Fig. 2d), which was accompanied by significant decreases in serum BUN and SCr compared to DMSO-treated IR controls, reflecting preserved renal function (Fig. 2e,f). Consistent with these protective effects, transcriptional profiling showed that ISRIB significantly attenuated the IR-induced increase in ISR-related gene expression (Fig. 2g) without impacting eIF2α phosphorylation (Extended Fig. 2c). Histological assessment of H&E-stained kidney sections demonstrated maintenance of tubular structure in ISRIB-treated animals compared to the IR group, which was further validated by quantitative analysis (Fig. 2h,i). Immunohistochemical analysis for KIM-1 showed further evidence of diminished tubular injury with ISRIB treatment (Fig. 2h,j). Morphological evaluation with AQP-1 also indicated better preservation of proximal tubular integrity in ISRIB-treated kidneys (Extended Fig. 2d). Together, these findings establish that ISR pathway activation exacerbates IR-induced renal injury, and that pharmacological blockade of ISR signaling provides substantial protection to renal proximal tubular cells against IR challenge.

### PKR is a critical enzyme that transmits mitochondrial stress signals to activate the ISR following renal IR injury

The identification of ISR involvement in IR-induced renal injury led us to explore the molecular factors responsible for mediating retrograde signaling from mitochondria to the nucleus. Recent evidence, including our own data, demonstrates that mitochondrial stress activates the ISR predominantly through Protein Kinase R (PKR), which acts as a key eIF2α kinase in this setting^40,41,51^. These findings suggest a central function for PKR in detecting mitochondrial dysfunction and facilitating signal transduction to the nucleus, thereby initiating ISR. Thus, we proposed that PKR is responsible for mediating mitochondrial dysfunction–induced ISR activation following renal IR injury.

To examine whether PKR mediates ISR activation during renal IR injury, we utilized a murine model with genetic deletion of *Pkr* (*Pkr⁻/⁻*)^41^. Both wild-type (*Pkr⁺/⁺*) and *Pkr⁻/⁻* mice underwent the established renal IR injury protocol (Fig. 3a). Gross anatomical assessment showed less severe renal damage in *Pkr⁻/⁻* mice compared to *Pkr⁺/⁺* mice (Fig. 3b). Furthermore, *Pkr⁻/⁻* mice maintained superior renal function, as reflected by decreased serum BUN and SCr values after IR injury (Fig. 3c,d). In agreement, microscopic evaluation of H&E-stained renal tissue revealed markedly diminished tubular injury in *Pkr⁻/⁻* mice (Fig. 3e,f). Additional immunohistochemical analysis for KIM-1 substantiated lower degrees of renal injury in *Pkr⁻/⁻* animals, further supporting the protective effect of PKR deficiency (Fig. 3e,g). The expected IR-induced increase in ISR-related gene expression seen in *Pkr⁺/⁺* mice was notably reduced in *Pkr⁻/⁻* mice, which corresponded with decreased phosphorylation of eIF2α (Fig. 3h,i). Taken together, these results establish PKR as a crucial mediator that connects mitochondrial stress and ISR activation during IR-induced renal injury.

**Fig. 3:**
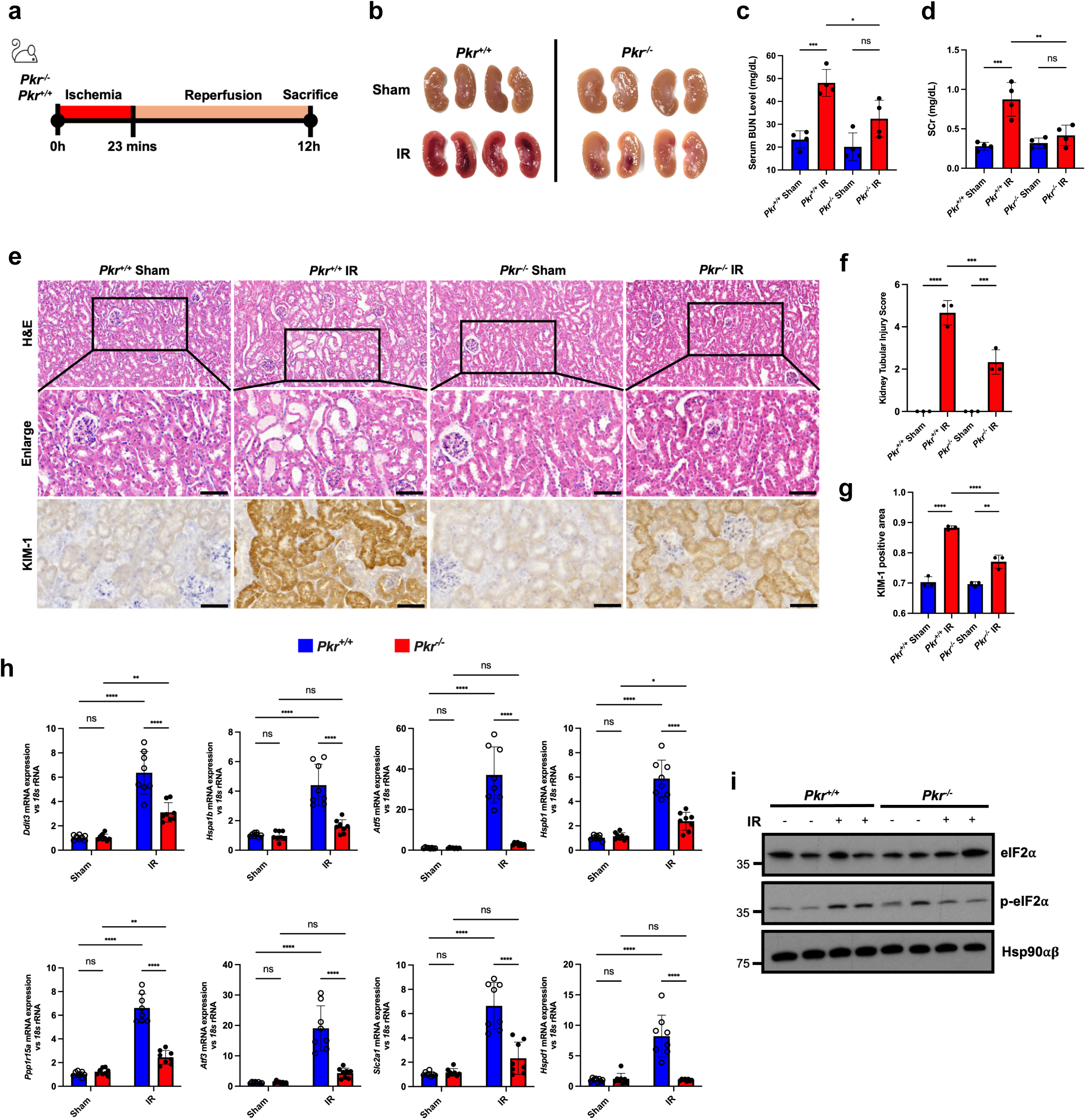
PKR is a critical mediator transmitting mitochondrial stress to activate ISR in renal IR injury. **a,** Schematic diagram illustrating bilateral IR-induced kidney injury in genetically modified mouse models *Pkr^+/+^* and *Pkr^−/−^*. **b,** Gross morphology of sham-operated and IR-injured kidneys in *Pkr^+/+^* and *Pkr^−/−^* mice. Photographs were taken immediately post-sacrifice following whole-body perfusion via the heart. **c-d,** Serum BUN and SCr levels were evaluated to assess kidney dysfunction after the IR procedure in *Pkr^+/+^* and *Pkr^−/−^* mice. *N* = 4. **e-g,** Representative H&E and KIM-1 immunohistochemistry staining of kidney sections from *Pkr^+/+^* and *Pkr^−/−^* mice. Tubular injury scores were determined by quantifying the percentage of tubules exhibiting injury-associated features as previously described. The KIM-1 positive area was quantified by normalizing chromogenic intensity to hematoxylin staining. *N =* 3. Scale bar = 30 µm. **h,** RT-qPCR analysis of ISR-related mRNA expression following IR in *Pkr^+/+^* and *Pkr^−/−^* mice demonstrates a significant downregulation of ISR-associated genes in *Pkr^−/−^* mice. *N =* 8. **i,** Immunoblot analysis showing eIF2⍺ and p-eIF2⍺ levels after kidney IR injury in *Pkr^+/+^* and *Pkr*^−/−^ mice. *Pkr^−/−^* mice displayed a significantly reduced level of p-eIF2⍺, a hallmark of ISR, supporting the role of PKR in regulating the ISR during IR-induced kidney damage. Hsp90 was used as the loading control. Data were analyzed by one-way ANOVA with multiple comparison correction unless otherwise specified. *\*P < 0.05, **P < 0.005, ***P < 0.001, ****P < 0.0001*.

### Kidney IR injury results in the accumulation of mitochondrial double-stranded RNAs

Given the central role of PKR in mediating IR-induced renal injury, we subsequently explored the molecular mechanism responsible for its activation. PKR is a double-stranded RNA (dsRNA)-dependent kinase whose activation is strongly associated with cellular stress responses^51–54^. Furthermore, prior research, including our own, has demonstrated that mitochondrial dysfunction results in the accumulation of mitochondrial dsRNAs, subsequently activating PKR signaling^41,55^. Given PKR’s involvement in IR-induced ISR activation, we proposed that mitochondrial stress during IR facilitates the accumulation of mitochondrial dsRNA, thereby inducing PKR activation. To validate this hypothesis, we analyzed cellular dsRNA levels using the J2 monoclonal antibody, which detects dsRNA in a sequence-independent manner^47^. Immunofluorescence analysis displayed a notable increase in J2 antibody staining in cells exposed to H/R, as well as in cells treated with GTPP, an established inducer of mitochondrial stress known to facilitate the release of mitochondrial dsRNA^41,56–58^, compared with control cells, indicating a significant accumulation of dsRNA under these stress conditions (Fig. 4a). Additionally, J2 antibody staining strongly co-localized with MitoTracker signals in H/R- and GTPP-treated cells^57,58^, suggesting that the accumulated dsRNA under these stress conditions is primarily of mitochondrial origin.

**Fig. 4:**
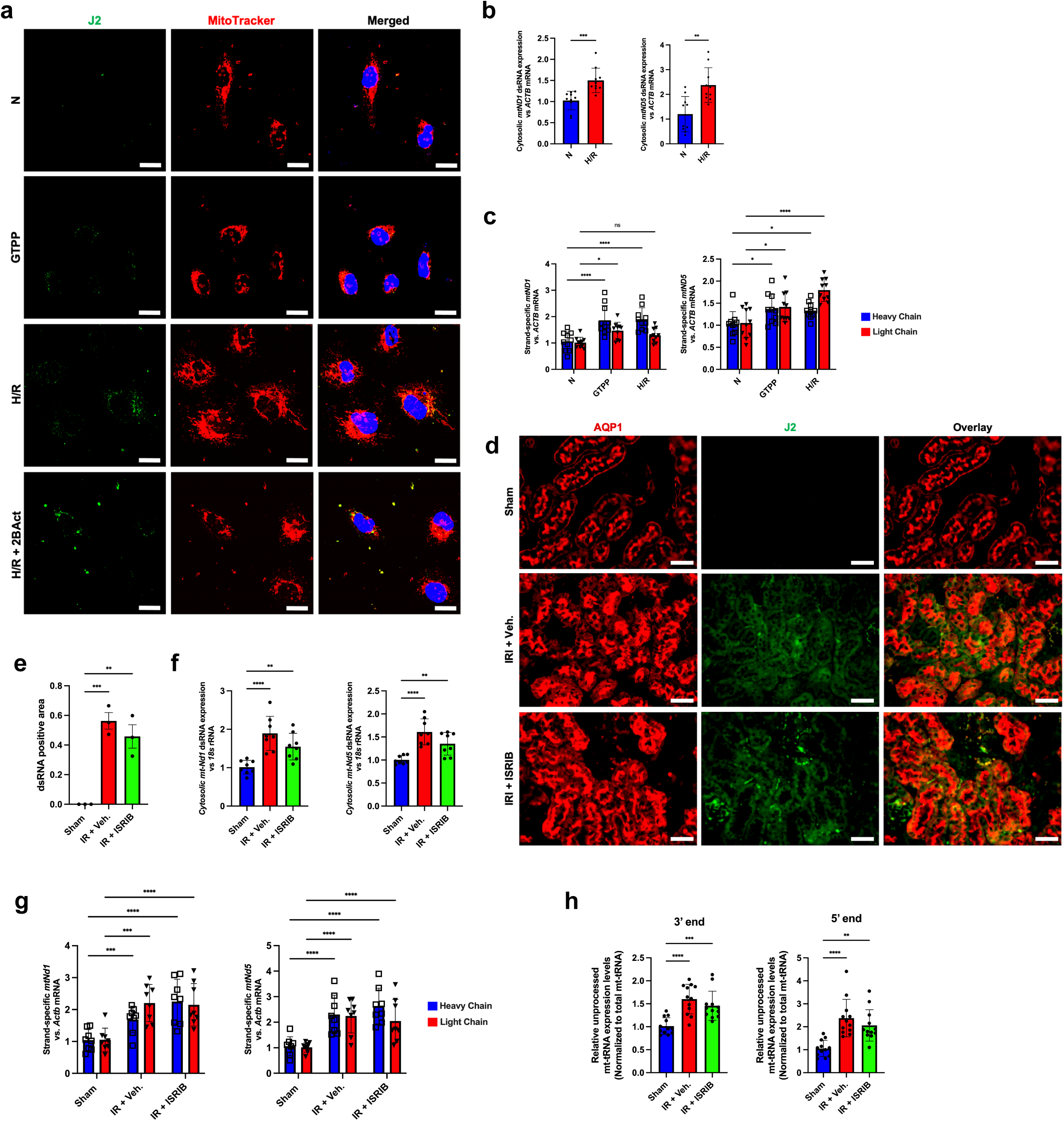
Kidney IR injury causes mitochondrial dsRNAs accumulation. **a,** Confocal imaging of J2-dsRNA immunostaining was performed on HK-2 cells under normoxic conditions and after 6 hours of hypoxia followed by 2 hours of reoxygenation. GTPP-treated HK-2 cells served as a positive control for evaluating dsRNA accumulation in response to mitochondrial stress. Mitochondrial localization of dsRNA was assessed using MitoTracker Orange CMTMRos as a mitochondrial marker. Scale bar = 20 µm. **b,** Gene expression analysis was conducted to evaluate cytosolic release of mitochondrial dsRNA, specifically assessing *mt-ND1* and *mt-ND5* levels in HK-2 cells following H/R, confirming the mitochondrial origin of dsRNA accumulation. *N =* 10. **c,** Cytosolic mitochondrial dsRNA was specifically detected in both heavy and light chains using strand-specific RT-qPCR in HK-2 cells subjected to H/R. GTPP treatment for 8 hours served as the positive control for evaluating cytosolic mitochondrial dsRNA release. *N =* 10. **d-e,** Immunostaining for J2-dsRNA in mouse kidney sections demonstrated substantial dsRNA accumulation in IR-induced kidney injury relative to sham controls. AQP1 staining was used to identify proximal tubule cells, distinguishing them from other renal cell types. Quantification of J2 intensity was done by normalizing to AQP1 intensity. *N =* 3. Scale bar = 50 µm. **f,** Assessment of cytosolic mt-Nd1 and mt-Nd5 gene expression in kidney tissue showed increased levels following IR and IR with ISRIB treatment, further supporting cytosolic release of mitochondrial dsRNA. *N =* 10. **g,** Cytosolic mitochondrial dsRNA specific detection for heavy and light chains was performed using strand-specific RT-qPCR in a mouse model of renal IR injury. *N =* 8. **h,** Mitochondrial RNA processing was analyzed by measuring levels of unprocessed mitochondrial tRNA^Met^ fragments, indicating defective mitochondrial RNA transcription, via RT-qPCR. *N =* 12. Statistical analysis employed a two-tailed unpaired t-test (**b,e,f,h**) and Tukey’s two-way ANOVA (**c,g**). *\*P < 0.05, **P < 0.005, ***P < 0.001, ****P < 0.0001*.

As PKR primarily resides in the cytosol, its activation requires the presence of dsRNA in this cellular compartment. Several studies have shown that mitochondrial stress leads to the production and accumulation of mitochondrial dsRNA, which can be transported into the cytosol via Bax/Bak channels, thereby activating cytosolic PKR^41,52,59^. Building on these findings, we quantified cytosolic mitochondrial dsRNA levels to assess whether H/R–induced mitochondrial stress facilitates dsRNA release into the cytosol. We measured cytosolic mitochondrial RNA content by RT–qPCR, which showed a substantial elevation of *mt-ND1* and *mt-ND5* transcripts in the cytosolic fraction of H/R-treated cells when compared to controls (Fig. 4b). Given that the mitochondrial genome is transcribed in both directions, we applied a strand-specific RT-qPCR approach to separately quantify sense and antisense mitochondrial transcripts, which allowed rigorous assessment of each strand and confirmation that PKR activation results from authentic mitochondrial dsRNA accumulation, since measurement of total cytosolic mitochondrial RNA alone does not discriminate between single-stranded and double-stranded forms^56,60^. The strand-specific qRT-PCR analysis indicated that both heavy- and light-strand mitochondrial transcript levels were significantly increased after GTPP and H/R treatments, supporting that mitochondrial dsRNA formation did indeed occur (Fig. 4c).

To translate our *in vitro* findings into an *in vivo* context, we conducted J2 immunofluorescence staining on mouse kidney sections to detect dsRNAs. After IR, renal tissues showed a marked increase in J2 fluorescence intensity, especially in renal proximal tubular epithelial cells, indicating heightened dsRNA accumulation that aligns with our *in vitro* findings (Fig. 4d,e). Having established cytosolic accumulation of mitochondrial dsRNAs under H/R stress *in vitro*, we next investigated whether this phenomenon also occurs *in vivo*. In mouse kidneys subjected to IR, mitochondrial RNAs were identified in the cytosolic fraction, providing evidence that mitochondrial dsRNA release may serve as a key factor in PKR activation and ISR induction (Fig. 4f). Furthermore, strand-specific analyses revealed increased levels of both heavy-and light-strand mitochondrial transcripts *in vivo*, mirroring the mitochondrial dsRNA accumulation and cytoplasmic release observed under H/R conditions in HK-2 cells (Fig. 4g).

Prior research has shown that mitochondrial dsRNA accumulation may result from defects in mitochondrial RNA processing^59^. To assess whether a similar impairment is present in our model, we evaluated mitochondrial RNA processing using primers for mitochondrial tRNA^Met^, which is typically cleaved from its polycistronic precursor^41,59^ (Extended Fig. 4a). Our analysis indicated altered mitochondrial RNA processing after IR, as evidenced by increased levels of unprocessed mitochondrial tRNA^Met^ with retention of both 3’ and 5’ flanking sequences on the polycistronic transcript (Fig. 4h).

Taken together, these data indicate that mitochondrial stress resulting from H/R or IR impairs mitochondrial RNA processing, leading to accumulation of mitochondrial dsRNAs that are subsequently released into the cytosol, triggering PKR activation and ISR signaling. Importantly, ISR inhibition did not alter the accumulation or cytosolic release of mitochondrial dsRNAs (Fig. 4a,e), supporting the conclusion that dsRNA generation and cytosolic export occur upstream of PKR–eIF2α signaling.

### Mitochondrial stress and subsequent ISR-mediated ferroptosis contribute to renal IR injury

Building on our observation that the ISR plays a pivotal role in IR-induced renal injury, we proceeded to investigate the specific mechanisms through which the ISR contributes to tissue damage. Traditionally, IR-induced renal injury has been mainly attributed to apoptotic and necrotic cell death^17,18,61^. Nevertheless, recent studies indicate that ferroptosis is a significant contributor to IR-mediated renal injury^33,62–67^. Furthermore, emerging evidence has shown that activation of the ISR can induce ferroptotic cell death via an ATF4-dependent pathway^68–70^. Collectively, these findings led us to hypothesize that IR-induced ISR might mediate renal injury, at least in part, through the process of ferroptosis.

To determine whether ferroptosis is activated in response to IR stress, we initially examined mouse and human transcriptomic datasets. Violin plots illustrating ferroptosis gene expression demonstrated that the ferroptosis pathway is highly conserved in both mouse and human transcriptomic data (Fig. 5A). In agreement, we observed a reduction in the glutathione metabolism pathway in the *in vivo* mouse IR dataset (Extended Fig. 5a). Collectively, these results suggest that ferroptosis is a key mechanism mediating IR-induced renal injury. In addition, well-characterized ferroptosis-related genes, such as *Chac1, Hmox1, Ptgs2, Fth1, Slc40a1, Slc7a11,* and *Nrf2*^34,71–73^, displayed differential expression following IR exposure (Fig. 5b). Correlation analysis revealed that upregulation of these genes was positively correlated (Extended Fig. 5b). RT–qPCR validation demonstrated significant induction of these ferroptosis markers in HK-2 cells subjected to H/R treatment (Fig. 5c), whereas protein levels of GPX4, a principal regulator of ferroptosis resistance^74^, were markedly decreased (Fig. 5d). To strengthen the evidence for ferroptotic lipid injury, we measured lipid peroxidation using BODIPY 581/591 C11^75^. Both H/R treatment and Erastin, a widely recognized ferroptosis inducer^76^, increased the proportion of oxidized BODIPY-positive cells, as assessed by flow cytometry (Extended Fig. 5c). Likewise, either H/R or Erastin administration enhanced oxidized BODIPY fluorescence, and this effect was substantially mitigated by the ferroptosis inhibitor Ferrostatin-1 (Fer-1)^77^ (Fig. 5e). Collectively, these findings demonstrate that H/R induces ferroptotic lipid peroxidation in HK-2 cells.

**Fig. 5:**
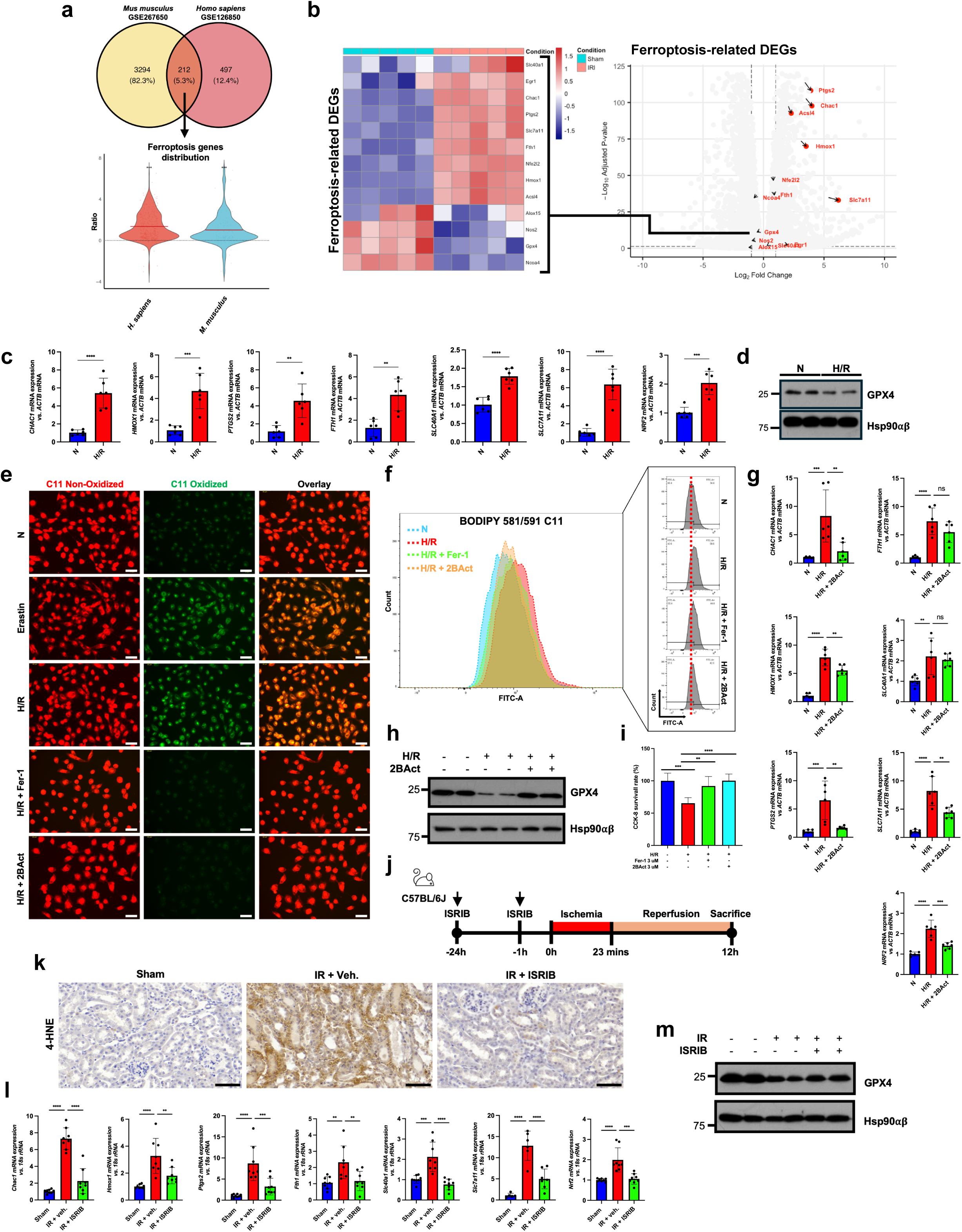
Mitochondrial stress and subsequent ISR-mediated ferroptosis is responsible for renal IR injury. **a,** Venn diagram (top) illustrating the overlap between differentially expressed genes from mouse kidneys subjected to 20 min ischemia and 16 h reperfusion (GSE267650) and paired human kidney biopsy transcriptomes collected before and after transplantation (GSE126805). From the shared gene pool, ferroptosis-associated transcripts were selected for visualization in the violin plot (bottom). Each dot represents the log_10_-normalized expression value of an individual ferroptosis gene; violin width indicates expression density and center lines denote the mean. **b,** Heatmap displaying DEGs linked to ferroptosis following IR-induced kidney injury compared to sham-treated mice, highlighting substantial upregulation of ferroptosis-associated genes. **c,** Gene expression analysis demonstrates changes in ferroptosis-related mRNA levels in HK-2 cells after 6 hours hypoxia followed by 2 hours reoxygenation. *N = 6.* **d,** Immunoblotting illustrates a reduced GPX4 level, indicating ongoing ferroptosis during the H/R protocol in HK-2 cells. **e,** BODIPY 581/591 C11 staining was performed on live HK-2 cells post-H/R, with or without co-treatment; Fer-1 was applied as a ferroptosis inhibitor during H/R, and 2BAct was used to inhibit ISR activation during H/R in HK-2 cells. As a ferroptosis-positive control, HK-2 cells treated with Erastin for 12 hours were included. Scale bar = 50 µM. **f,** Flow cytometry after BODIPY 581/591 C11 staining of live HK-2 cells post-H/R indicated that Fer-1 and 2BAct co-treatment led to reduced levels of oxidized C11, indicative of improved ferroptotic status with lower lipid peroxidation. **g,** Gene expression analysis showed that ISR inhibition via 2BAct co-treatment resulted in reduced expression of ferroptosis-related mRNAs in H/R-stimulated HK-2 cells. *N = 6.* **h,** Restoration of GPX4 protein was observed after ISR inhibition by 2BAct following H/R in HK-2 cells, as demonstrated by immunoblot; Hsp90 was utilized as the protein loading control. **i,** CCK-8 cell viability analysis following H/R in HK-2 cells revealed enhanced cell survival with ISR inhibition by 2BAct, in agreement with the inhibition of ferroptosis by Fer-1. *N =* 6. **j,** Schematic of the experimental protocol for bilateral IR-induced kidney injury in mice, incorporating ISRIB administration. **k,** 4-HNE-immunohistochemistry of kidney sections demonstrated smaller injury areas in mice pretreated with ISRIB before IR induction. Scale bar = 60 µm. **l,** RT-qPCR analysis showed decreased expression of ferroptosis-related mRNAs in IR injury kidneys from mice injected with ISRIB. *N =* 8. **m,** Immunoblot analysis revealed restoration of GPX4 levels in kidneys subjected to IR injury after ISRIB administration. Data were assessed using two-tailed unpaired t-test **(c, i)** and one-way ANOVA for multiple group comparisons unless otherwise stated. *\*P < 0.05, **P < 0.005, ***P < 0.001, ****P < 0.0001*.

We next examined whether this process is regulated by the ISR. Pharmacological inhibition of the ISR with 2BAct or inhibition of ferroptosis with Fer-1 significantly decreased H/R-induced lipid peroxidation, as indicated by lower oxidized BODIPY C11 fluorescence and a reduced population of oxidized cells in flow cytometric analyses (Fig. 5e,f). In addition, 2BAct notably diminished the H/R-induced upregulation of ferroptosis-related gene expression (Fig. 5g) and restored GPX4 protein expression that was otherwise suppressed by H/R (Fig. 5h). Consistently, cell viability assays showed that treatment with either Fer-1 or 2BAct substantially prevented H/R-induced cell death (Fig. 5i), supporting the role of ISR activation in mediating ferroptotic cell death under H/R conditions. Moreover, we observed that iron levels, a key indicator of ferroptosis^78^, were significantly reduced following 2BAct and Fer-1 treatment in H/R-challenged HK-2 cells, suggesting a protective effect against H/R-induced cellular damage (Extended Fig. 5d,e).

To further corroborate these results *in vivo*, we conducted a renal IR experiment as described previously (Fig. 5j). Immunostaining for 4-HNE, a marker of lipid peroxidation^34^, demonstrated a significant reduction in IR-induced lipid peroxidation in ISRIB-treated mice when compared to vehicle-treated controls (Fig. 5k). Similarly, BODIPY 581/591 C11 staining of freshly isolated kidney tissue showed a marked reduction in oxidized lipid content after ISRIB treatment (Extended Fig. 5f). At the transcriptional level, ISRIB administration lowered the IR-induced expression of genes associated with ferroptosis (Fig. 5l), while immunoblot analysis indicated restoration of GPX4 protein levels (Fig. 5m). Taken together, these findings indicate that ISR inhibition reduces ferroptosis *in vivo* during renal IR injury, highlighting an essential role for ISR signaling in the regulation of ferroptosis.

### PKR mediates ferroptosis during renal IR injury via the accumulation of mitochondrial dsRNA through activation of the ISR

Pharmacological inhibition of the ISR reduced ferroptosis during renal IR injury, leading us to examine the upstream activator of this pathway. In light of the protective function of PKR described above, we exposed *Pkr^⁻/⁻^* mice to IR injury (Fig. 6a). Aligning with our previous results, *Pkr^⁻/⁻^* mice exhibited protection from IR-induced renal injury, displaying significantly improved renal morphology and decreased tissue damage compared to wild-type controls (Fig. 6b). Likewise, *Pkr^⁻/⁻^* kidneys showed notably diminished lipid peroxidation, demonstrated by reduced 4-HNE staining (Fig. 6c). This decrease in lipid peroxidation was paralleled by downregulation of ferroptosis-associated genes (Fig. 6d) and maintenance of GPX4 protein expression (Fig. 6e), collectively supporting a reduction of ferroptotic cell death in the absence of PKR.

**Fig. 6:**
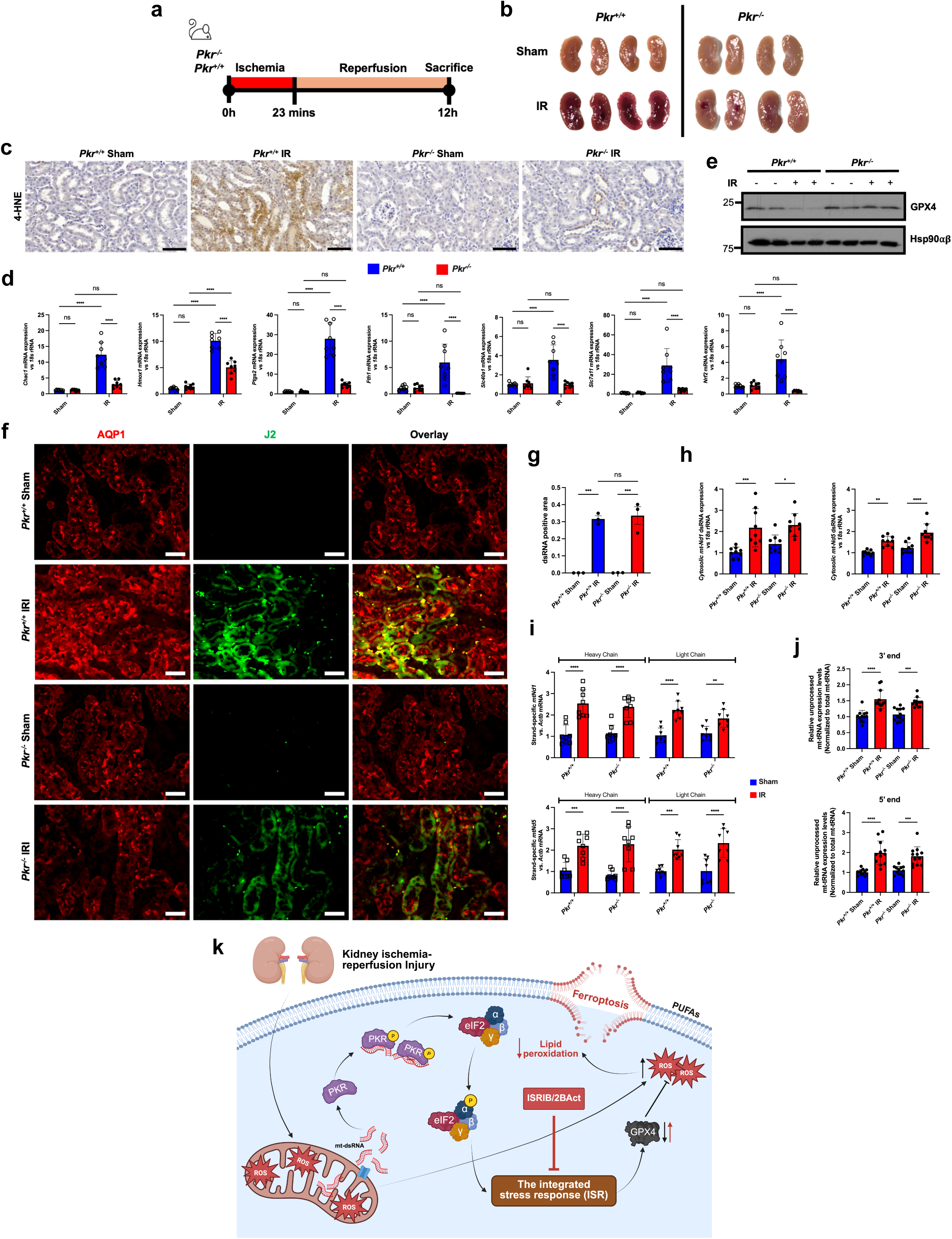
PKR mediates ferroptosis during renal IR injury via accumulation of mitochondrial dsRNA through the ISR. **a,** Experimental design for the induction of bilateral IR kidney injury in genetically modified mice (*Pkr^+/+^* and *Pkr^−/−^*). **b,** Gross morphological appearance of kidneys from sham-treated and IR-injured *Pkr^+/+^* and *Pkr^−/−^* mice. **c,** Immunohistochemical staining for 4-HNE demonstrates reduced lipid peroxidation in the kidneys of IR-injured *Pkr^−/−^* mice. Scale bar = 60 µm. **d,** RT-qPCR analysis of ferroptosis-associated mRNA reveals decreased expression of ferroptotic genes in kidneys from IR-injured *Pkr^−/−^* mice. *N =* 8. **e,** Immunoblot analysis of GPX4 indicates higher protein levels in IR-injured *Pkr^−/−^* kidneys compared to *Pkr^+/+^*, reflecting reduced activation of ferroptotic processes following IR injury. Hsp90 is used as a protein loading control. **f-g,** J2 antibody immunostaining of proximal tubule cells to visualize dsRNA in IR-injured *Pkr^+/+^* and *Pkr^−/−^* mouse kidneys, in comparison to sham controls. AQP1 staining marks tubular cells for precise identification among kidney cell populations. Quantification of J2 intensity is normalized to AQP1 intensity. *N =* 3. Scale bar = 50 µm. **h,** RT-qPCR analysis demonstrates similar levels of cytosolic *mt-Nd1* and *mt-Nd5* transcripts between IR-injured *Pkr^+/+^* and *Pkr^−/−^* kidneys. *N =* 9. **i,** Analysis of cytosolic mitochondrial dsRNA using strand-specific RT-qPCR reveals comparable expression of heavy and light mitochondrial dsRNA strands in renal IR injury between *Pkr^+/+^* and *Pkr^−/−^* mice. *N =* 8. **j,** Assessment of mitochondrial RNA processing via quantification of unprocessed mitochondrial tRNA^Met^ fragments by RT-qPCR shows no significant difference in unprocessed tRNA levels, indicating equivalent mitochondrial RNA processing impairment between *Pkr^+/+^* and *Pkr^−/−^* mice after renal IR INJURY. *N =* 12. **k,** Diagram illustrating the proposed signaling mechanism: mitochondrial dysfunction during renal IR injury results in the accumulation and cytosolic release of mitochondrial dsRNA, which is recognized by PKR. PKR subsequently phosphorylates eIF2⍺, serving as a central regulator of ISR activation, thereby enhancing lipid peroxidation and ferroptosis. ISRIB functions to inhibit this pathway during renal IR injury, consequently reducing lipid peroxidation and limiting ferroptotic damage. Created with Biorender.com. Publication license obtained. Data were analyzed using Tukey’s two-way ANOVA (**i**), two-tailed unpaired t-test **(g),** and one-way ANOVA with multiple comparisons unless indicated otherwise. *\*P < 0.05, **P < 0.005, ***P < 0.001, ****P < 0.0001*.

Immunofluorescence analysis using the J2 antibody demonstrated prominent mitochondrial dsRNA accumulation in both *Pkr^+/+^* and *Pkr^⁻/⁻^* kidneys following IR (Fig. 6f,g). Comprehensive quantitative and strand-specific RT–qPCR analysis revealed similar levels of cytosolic mitochondrial dsRNA fractions (Fig. 6h,i) and indicated unchanged mitochondrial RNA processing between genotypes (Fig. 6j), suggesting that PKR does not regulate mitochondrial dsRNA generation. Altogether, these findings support a model where mitochondrial dsRNA accumulation during IR activates PKR, triggering the ISR and promoting ferroptosis (Fig. 6k). The absence of PKR prevents ISR activation, thereby reducing ferroptotic damage and safeguarding renal structure. As such, PKR serves as an essential molecular link between mitochondrial RNA stress and ISR-mediated ferroptosis in renal IR injury.

## Discussion

Renal IR injury is a leading cause of acute kidney injury (AKI) and is associated with severe mitochondrial dysfunction and oxidative stress^9,17,29,35^. Despite extensive research, the molecular pathways that connect mitochondrial disruption to stress response signaling and cell fate decisions remain not fully elucidated. In this study, we uncover a previously uncharacterized pathway in which mitochondrial dysfunction leads to dsRNA accumulation in mitochondria, initiating PKR-mediated ISR signaling that subsequently induces ferroptotic cell death after renal IR injury. These results provide mechanistic insights into the interplay between mitochondrial stress and regulated cell death, unveiling a new mitochondrial-to-nuclear signaling and ferroptosis axis that drives IR-induced kidney injury.

Consistent with previous reports, which identify mitochondrial perturbation as an early hallmark of IR injury^9,17,29,35^, transcriptomic and histological analyses across mouse, human, and cell models revealed pronounced mitochondrial stress. These IR-induced metabolic disturbances were associated with strong transcriptional upregulation of ISR target genes and greater phosphorylation of eIF2α, indicating robust ISR activation. Although the ISR is classically viewed as a cytoprotective mechanism that restores proteostasis^79–81^, our results indicate that excessive or prolonged ISR activation exacerbates renal injury under ischemic-reperfusion stress, consistent with prior evidence that strong ISR activation can increase damage potential^81–83^. Pharmacological inhibition of the ISR by ISRIB or 2BAct, both of which are selective eIF2B activators, significantly enhanced cell viability and protected renal function both *in vitro* and *in vivo*, demonstrating a causal contribution of maladaptive ISR signaling to IR-induced tissue injury. Of the four canonical ISR kinases—PERK, HRI, GCN2, and PKR^81^—our findings identify PKR as the main mediator responsible for ISR activation in renal IR injury. Genetic ablation of PKR reduced eIF2α phosphorylation, suppressed ISR gene expression, and mitigated tissue injury, further supporting a critical role for PKR in this maladaptive pathway.

During ischemia, oxygen deprivation collapses oxidative phosphorylation and the tricarboxylic acid (TCA) cycle, compelling mitochondria in renal cells to switch to anaerobic metabolism^22,84^. Upon reperfusion, the sudden restoration of oxygen induces excessive electron leakage from the over-reduced electron transport chain, resulting in an overproduction of ROS and further mitochondrial injury^18,85^. This oxidative stress compromises mitochondrial integrity and interferes with bidirectional transcription of the circular polycistronic mitochondrial genome^55^, ultimately disrupting mitochondrial RNA metabolism. Consistently, our findings demonstrated that mitochondrial dysfunction induced by IR stress impairs mitochondrial RNA processing, resulting in the accumulation of aberrantly processed transcripts that anneal to form mitochondrial dsRNA. Under physiological condition, both mitochondrial DNA strands are co-transcribed to generate sense and antisense RNAs, which undergo precise processing and are eliminated by mitochondrial RNA surveillance mechanisms^60,86–88^. However, in the context of IR injury, mitochondrial damage undermines this quality-control system, leading to mitochondrial dsRNA accumulation and its leakage into the cytosol^86,89^. Once present in the cytosol, mitochondrial dsRNA interacts with PKR, triggering downstream ISR signaling. This model is supported by emerging evidence that failure of mitochondrial RNA surveillance produces cytosolic dsRNA species that activate antiviral-like stress responses^51,52,86^. Notably, inhibition of ISR did not affect the accumulation of mitochondrial dsRNA, suggesting that mitochondrial RNA dysregulation precedes PKR activation and the subsequent engagement of the ISR.

Ferroptosis has been increasingly implicated as a contributor to IR-induced renal injury^28,33,64,90,91^. In agreement with these findings, our results further demonstrate that IR triggers ferroptotic cell death, indicated by a reduction in GPX4 expression and an elevation in lipid peroxidation, supporting the view that ferroptosis is a critical mediator of kidney injury. Mitochondrial dysfunction during IR appears to have a pivotal role in initiating ferroptosis, as oxygen deprivation during ischemia impairs oxidative phosphorylation, whereas reperfusion generates a significant increase in ROS that oxidizes polyunsaturated fatty acids in cellular membranes, ultimately disrupting membrane stability^18,33,92,93^. Under normal physiological conditions, GPX4 counters lipid hydroperoxides by utilizing GSH as a reductant to uphold redox equilibrium^94,95^. Conversely, mitochondrial damage decreases NADPH generation, hampers the regeneration of GSH, and exceeds the detoxification ability of GPX4, thereby driving enhanced lipid peroxidation and ferroptosis^96,97^. Concurrently, we identified strong ISR activation in response to IR injury. While transient ISR signaling confers cytoprotection, sustained ISR marked by persistent eIF2α phosphorylation promotes oxidative stress and ferroptotic cell death^42,98^. Our earlier research further established that ATF4, a key ISR effector, elevates REDD1, thereby suppressing mTOR signaling and promoting GPX4 degradation, which in turn amplifies ferroptosis and cell death^99^. Other study has also revealed that CHAC1, a critical glutathione metabolism regulator, was a direct target of the ATF4 as an ISR effector and regulate pro-ferroptotic response and the ablation of CHAC1-ATF4 axis mediated a protective effect against ferroptosis.^73^ Consistent with this mechanism, IR-induced activation of the ISR was associated with decreased GPX4 levels and increased ferroptotic signaling during kidney injury. Notably, pharmacological inhibition of the ISR using selective eIF2B activators restored GPX4 expression, reduced lipid peroxidation, and preserved renal function, indicating that ISR hyperactivation promotes ferroptosis. Further supporting this mechanism, genetic deletion of PKR—the principal ISR kinase activated in our model—reduced eIF2α phosphorylation, diminished ferroptotic signaling, and alleviated renal injury. Together, these findings establish ISR activation, particularly via PKR, as a key regulator of ferroptosis and a significant factor in IR-induced kidney damage.

Although our new findings advance our understanding of mitochondrial dsRNA–mediated ISR activation in IR-induced ferroptosis, several important issues remain unresolved. The precise mitochondrial defects that result in impaired RNA processing and mitochondrial dsRNA accumulation have yet to be fully characterized. Previous research demonstrated that deficiencies in PNPase, an RNA helicase involved in mitochondrial RNA processing, led to worsened mitochondrial dysfunction and dsRNA buildup following IR^47,88^, indicating that PNPase may represent a potential mechanistic contributor to our observations. Subsequent investigations should clarify the roles of mitochondrial transcription and RNA degradation systems, including SUV3 and mitochondrial RNA granule-associated proteins^88,100,101^. While PKR is identified as the principal ISR kinase activated in renal IR injury, the involvement of additional eIF2α kinases, such as PERK, GCN2, and HRI, should be further investigated. Additionally, considering the major function of iron metabolism in ferroptosis, it is critical to evaluate how ISR activation affects iron homeostasis, including regulation of ferritin, transferrin receptor, and iron–sulfur cluster biosynthesis under IR-induced stress.

In summary, this study identifies a novel pathway that connects mitochondrial dysfunction with ferroptotic cell death in renal IR injury. Our data establish mitochondrial dsRNA accumulation as an intrinsic stimulus of the IR-induced ISR through PKR activation, thus expanding the known physiological roles of mitochondrial dsRNA from antiviral and inflammatory responses to include sterile tissue injury. Moreover, we delineate a mechanistic association between ISR activation and ferroptosis, demonstrating that ISR hyperactivation following IR enhances ferroptotic death and aggravates renal injury, while pharmacological ISR inhibition mitigates ferroptosis and supports renal function. Together, these results define the mitochondrial dsRNA–PKR–ISR pathway as a central regulator of IR-induced ferroptosis and position ISR modulation as a compelling therapeutic target for acute kidney injury, mitochondrial stress–related conditions, and other IR-associated disorders.

## Acknowledgements

We thank H.Y. Kwon (Soonchunhyang Institute of Medi-bio Science) for the provision of HK-2 cells utilized in *in-vitro* experiments, and Y.S. Hwang (Soonchunhyang Institute of Medi-bio Science) for supplying hypoxia-related equipment. Cell imaging and flow cytometry were conducted using the Zeiss LSM710 confocal microscope and the BD FACS Canto II flow cytometer at the Soonchunhyang Biomedical Science Core-facility of the Korea Basic Science Institute (KBSI). This study received support from the Global-Learning & Academic research institution for Master’s·PhD students, and Postdocs (G-LAMP) Program of the National Research Foundation of Korea (NRF) grant (No. RS-2025-25441283), as well as the NRF grant (NRF-2022R1A2C1010676) funded by the Ministry of Education.

## Contributions

R.J. and J.H. initiated the study, devised the experimental design, and drafted the manuscript. R.J. was primarily responsible for the mouse and *in vitro* experiments. M.S.C., S.Y.P., N.L., and Y.Y. assisted with experiments related to the mouse kidney IR model. R.J. and J.K. undertook the bioinformatics analyses. H.G. and S.P. participated in the interpretation and analysis of mouse experimental data. F.K. assisted with mitochondrial processing analysis. K.A.N. provided technical support and contributed to experimental planning. All authors engaged in discussions and gave approval to the final manuscript.

## Methods

### Chemicals and reagents

The chemicals utilized in this study include Erastin (HY-15763, MedChemExpress, USA), ISRIB (SML0843, Sigma-Aldrich, USA), Ferrostatin-1 (F1302, Tokyo Chemical Industry, Japan), and Gamitrinib-triphenylphosphonium (GTPP), which was obtained from LegoBiosciences Inc. (Daejeon, Republic of Korea) as previously described^102^. 2BAct was synthesized by the Medicinal Chemistry Team at Dageu-Gyeongbuk Medical Innovation Foundation (Daegu, Republic of Korea)^99^. Additional reagents included DAB substrate (SK-4100, Vector Laboratories, USA), ClearView^TM^ hematoxylin (MA0101010, StatLab, USA), ClearView^TM^ eosin (MA0101015, StatLab, USA), cell-counting kit 8 (CCK-8) (CK04, Dojindo, Japan), FerroOrange (F374, Dojindo, Japan), creatinine assay kit (ab65340, Abcam, UK), QuantiChrom^TM^ urea assay kit (DIUR-100, BioAssay Systems, USA), ReverTra Ace RT master mix (TOFSQ-201, Toyobo, Japan), SYBR green (RT501M, Enzynomics, Republic of Korea), protease inhibitor cocktail (P3100-005, GenDEPOT, USA), phosphatase inhibitor cocktail (P3200-001, GenDEPOT, USA), Pierce^TM^ BCA protein assay kit (23225, Thermo Fisher Scientific, USA), PVDF membrane (IPVH00010, Merck Millipore, USA), ClaroSola enhanced chemiluminescent (ECL) solution (HQS071, BioD, Republic of Korea), MitoTracker^TM^ Orange CMTMRos (M7510, Invitrogen, USA), Hoechst 33342 Trihydrochloride Trihydrate (H3750, Invitrogen, USA), SuperScript^TM^ IV Reverse Transcriptase (18090050, Invitrogen, USA), and BODIPY 581/591 C11 (D3861, Invitrogen, USA).

### Murine and experimental models of renal IR injury

The animal experimental procedure and protocol were approved by the Soonchunhyang University Animal Care and Use Committee (SCH24-0002) and were rigorously performed in compliance with the established guidelines. Mice were anesthetized throughout the experimental procedure and during blood-tissue collection, which was followed by euthanasia using cervical dislocation. All mice were housed in a specific-pathogen-free (SPF) facility with a 12-h light/dark cycle at a constant temperature of 22±1 °C, relative humidity of 30%-70%, and had unlimited access to lab diet (PicoLab Mouse Diet 20-5058, Lab Diet). *Pkr^−/−^* (knockout) mice were procured from the Australian Phenomics Facility (APF, APB631) and were backcrossed onto the C57BL/6J background for over six generations. In brief, 8-9-week-old mice were anesthetized using isoflurane administered via animal inhalation narcosis device (JD-C-107A, JEUNGDO Bio & Plant CO., LTD, Republic of Korea). To induce IR injury, a midline laparotomy was carried out, and both renal pedicles were occluded with non-traumatic serration micro vascular clamps (RS-5424, Roboz) for 23 mins, while body temperature was maintained at 34-37 °C using a temperature-controlled surgical mattress. Control or sham mice underwent identical procedures without the clamping of the kidneys. Upon completion of the ischemic interval, reperfusion was verified visually at clamp removal by observing the color change from purplish to the normal kidney appearance. To inhibit ISR activation during IR, 5 mg/kg of ISRIB was administered to mice at 24 h and 1 h before the surgery. After the IR procedure, the incision was closed, followed by subcutaneous saline injection, and the mice were placed under a heating lamp for recovery. Mice were euthanized 12 hours after recovery, and then blood and tissue samples were collected.

### Cell culture, treatment, and hypoxia/reoxygenation conditioning

Normal human kidney proximal tubular epithelial cells (HK-2, kindly provided by H.Y. Kwon, Soonchunhyang Institute of Medi-bio Science) were maintained in DMEM/F12 (Gibco) supplemented with 10% fetal bovine serum (FBS, Gibco) and 1% penicillin/streptomycin (Corning) in a humidified incubator at 5% CO_2_ and 37 °C. To simulate IR injury, HK-2 cells were subjected to H/R by incubating for 6 hours in a hypoxic environment (1% O_2_, 5% CO_2_, and 94% N_2_) using serum- and glucose-free media, which had been saturated overnight prior to media exchange in a hypoxia chamber (InvivO_2_, Baker Co.), followed by reoxygenation by replacing with standard media and returning cells to a normoxic incubator for the designated recovery time before harvesting. To assess the effect of ISR inhibition during H/R, HK-2 cells received pre-treatment with 3 μM 2BAct and 3 μM ISRIB for 1 h before hypoxic exposure and were exposed to the same treatments during the reoxygenation phase. For induction of ferroptosis and mitochondrial stress to promote mitochondrial dsRNA release, HK-2 cells were treated with 10 μM Erastin and 20 μM GTPP, respectively.

### Histopathological and immunofluorescence analysis

Mouse kidney tissues were fixed with 4% paraformaldehyde (SM-P01-100, GeneAll, Korea) and processed with ethanol prior to paraffin embedding (39601095, Leica) according to standard protocols. In brief, 5-μm tissue sections were deparaffinized in xylene and subsequently rehydrated through a descending ethanol concentration series. Endogenous peroxidase activity was quenched using 3% H_2_O_2_ (081-04215, Fujifilm), and antigen retrieval was performed by heating the sections in 1 mM EDTA at 95 °C using a high-pressure cooker for 20 mins, followed by cooling at room temperature for 20 mins. For permeabilization, the sections were incubated at room temperature for 15 mins with 0.1% Triton-X in PBS and then blocked for 30 mins in 5% BSA in PBS. Primary antibody incubation (see Supplementary Table 1) was performed overnight at 4 °C, and this was followed by a 2-hour incubation with HRP-conjugated secondary antibody. Diaminobenzidine (DAB) substrate was used for chromogenic staining, followed by counterstaining with hematoxylin. Samples were then visualized using a microscope (BX51, Olympus) and scanned with a slide scanner (MoticEasyScan, Motic). For immunofluorescence analysis, the sections were incubated for 2 hours with fluorescence-conjugated secondary antibody. Fluorescent images were acquired using a fluorescence microscope (DMi8, Leica). Quantification of DAB-positive areas and fluorescence intensity was performed using the color deconvolution plugin in ImageJ 1.54g software. For H&E staining, sections were mounted with DPX mounting medium (06522, Sigma-Aldrich) and renal tubular injury scores were obtained by assessing the percentage of tubules showing features such as vacuolization, necrosis, desquamation, dilatation, atrophy, cast formation, edema, and immune cell infiltration. Scoring was performed via randomized-blind analysis and classified as follows; none (0), 1-10% (1), 11-25% (2), 26-50% (3), 51-75% (4), and 76-100% (5).

### Cell viability assay

To assess cell viability following H/R and treatment, HK-2 cells were seeded at a density of 10,000 cells per well in a 96-well plate and incubated overnight. For reference, cells in the control group were pre-treated with Fer-1 for 1 hour before H/R and maintained with Fer-1 during reoxygenation. After completing the H/R procedure, CCK-8 reagent was added according to the manufacturer’s instructions and incubated for 1 hour. Absorbance was then measured at 450 nm with a microplate reader (Multiskan GO, Thermo Fisher Scientific).

### Iron measurement assay

To evaluate intracellular iron levels following H/R induction, HK-2 cells were stained with FerroOrange. After H/R treatment, cells were washed with 1X HBSS and stained with 1μM FerroOrange in HBSS for 30 mins. Fluorescence was measured at 543 nm excitation and 580 nm emission using a luminometer (GloMax Discover, Promega) and visualized with a confocal microscope (LSM710, Zeiss) at the same excitation and emission settings.

### Kidney function assessment assay

Serum creatinine (SCr) and blood urea nitrogen (BUN) assays were performed to evaluate kidney function following the IR procedure. Briefly, blood samples were collected from mice via the inferior vena cava at the time of sacrifice, and blood components were separated by centrifugation. Plasma was isolated and analyzed in accordance with each kit’s instructions.

### Real-time quantitative polymerase chain reaction (RT-qPCR)

RNA was isolated from bead-homogenized kidney or cells using Trizol by phase separation and RNA precipitation, following standard protocol. A two-step RT-qPCR was conducted to assess mRNA levels of target genes. Briefly, 1 μg of total RNA was reverse-transcribed into cDNA using a cDNA synthesis kit according to the manufacturer’s instructions. The resulting cDNA was then subjected to qPCR employing SYBR Green master mix and analyzed with the QuantStudio 1 Real-Time PCR system (Applied Biosystems). Data were normalized and processed using the 2^−ΔΔCt^ method. Primer sequences are provided in Supplementary Table 2.

### Western blot analysis

Protein extraction was performed on bead-homogenized kidney or cells using RIPA buffer (25 mM Tris-HCl pH 7.6, 150 mM NaCl, 1% NP-40, 1% sodium deoxycholate, and 0.1% SDS) supplemented with protease and phosphatase inhibitor cocktail. Purified proteins were obtained through mechanical and chemical lysis, and protein concentration was determined using the BCA Protein Assay Kit. Proteins were separated by SDS-PAGE based on molecular weight and transferred to polyvinylidene fluoride (PVDF) membranes for detection of target proteins. Membranes were blocked with 5% skim milk in 1X TBS-T, incubated overnight at 4 °C with primary antibodies, followed by a 2-hour incubation with HRP-conjugated secondary antibodies at room temperature. Each incubation step was followed by three washes with 1X TBS-T. Protein expression was visualized using enhanced chemiluminescence (ECL) substrate per the manufacturer’s instructions, and detected via X-ray film exposure with an automatic film processor (CP1000, Agfa) in a darkroom.

### Mitochondrial dsRNA localization microscopy

To monitor mitochondrial stress-induced dsRNA accumulation, HK-2 cells were treated with 20 μM GTPP for 8 hours and subjected to H/R as previously described. Mitochondria were labeled with MitoTracker 30 mins before the end of the procedure. In summary, cells were washed with 1X PBS and fixed using 4% PFA on a confocal-specialized cell culture dish (SPL) for 15 mins at room temperature, followed by three washes with 1X PBS. To permeabilize the cells, 0.1% v/v Triton X-100 in PBS was applied for 10 mins, then blocked with 5% BSA in PBS for 45 mins at room temperature. Subsequently, cells were incubated with J2 antibody at 4 °C overnight, washed three times with 1X PBS, and incubated with a fluorescence-conjugated secondary antibody at room temperature for 2 hours. Hoechst was then added to stain the nuclei, and samples were imaged using a confocal microscope (LSM710, Zeiss) and analyzed with Zeiss ZEN 3.9 software.

### Detection of mitochondrial dsRNAs accumulation

Total mitochondrial dsRNAs corresponding to *mt-ND1* and *mt-ND5* were measured by RT-qPCR using primers specific for transcripts derived from the polycistronic mt-DNA. For analysis of double-stranded RNA formation, HK-2 cells underwent strand-specific assessment, in which heavy and light chain mt-RNA transcripts were separated according to a previously established protocol with minor modifications^56^. In brief, total RNA was isolated using the Trizol-acid phenol-chloroform technique, followed by isopropanol precipitation. Target RNAs were then reverse transcribed into cDNA using SuperScript^TM^ IV Reverse Transcriptase (Invitrogen) with primers incorporating a CMV tag to facilitate strand amplification. The resulting cDNAs, representing specific mt-RNAs, were quantified via RT-qPCR and normalized to β-actin mRNA levels. All primer sequences are provided in Supplementary Table 3.

### RNA processing level analysis

To investigate mitochondrial function at the level of RNA processing, total RNAs from mouse samples were isolated, followed by cDNA synthesis and RT-qPCR analysis as previously described. Unprocessed tRNA^Met^ generated during mitochondrial RNA processing was utilized as an indicator of RNA processing failure, as outlined in earlier studies^41^. In summary, primers targeting the RNase P cleavage site at the 5’ end and an internal region of tRNA^Met^ were designed to assess failed processing upstream of the RNA transcript. Correspondingly, primers designed for the RNase Z cleavage site at the 3’ end and an internal region of tRNA^Met^ were employed to evaluate failed processing downstream of the RNA transcript. A detailed schematic of the mechanism is provided in Supplementary Figure 4. The abundance of each sequence was normalized using the Ct values from primers targeting the internal region of tRNA^Met^, thereby representing the total tRNA^Met^ in the total RNA. The sequences and primer information used in this study are detailed in Supplementary Table 4.

### Flow cytometry

For the assessment of ferroptosis, HK-2 cells underwent H/R protocol as previously described. HK-2 cells treated with 10 μM Erastin served as the positive control for ferroptosis. In brief, cells were stained with BODIPY 581/591 C11 mixed into growth media for 30 mins in the cell incubator. The culture medium was then collected, and cells were washed three times with pre-chilled 1X HBSS (Corning) and detached using Trypsin (Corning). All solutions were combined, and cells were pelleted by centrifugation. To reduce excessive staining, cells were rinsed with 1X HBSS and pelleted once more before being resuspended in 1X HBSS for flow cytometry analysis. The non-oxidized and oxidized forms of BODIPY 581/591 C11 were detected by measuring fluorescence emission using the BD FACSCanto^TM^ II Flow Cytometer System (Becton Dickinson, USA). Data analysis was conducted with FlowJo software (v10.10).

### Assessment of lipid peroxidation

Sections of kidney tissue (5 μm) prepared in optimal cutting temperature (OCT) compound (HI0-0051/#4583, Sakura) were rinsed in 1X PBS and stained with 2 μM BODIPY 581/591 C11 for 40 mins. Hoechst was then added for nuclear staining, and samples were visualized by fluorescence microscopy (DMi8, Leica). For HK-2 cells, BODIPY 581/591 C11 was administered 30 mins prior to completion of the treatment. After triple washing with 1X PBS, live cells were visualized using a fluorescence microscope (CELENA X, Logos Biosystems), followed by further analysis using CELENA X Cell Analyzer software.

### RNAseq analysis and data processing

Bioinformatics assessment was performed using the public dataset GSE267650, which includes RNAseq data from mice subjected to a 20-minute renal vascular clamp followed by 16 hours of reperfusion to model the IR procedure; gene expression in these samples was analyzed relative to sham-treated mice. For human samples, the public dataset GSE126805, consisting of patients undergoing kidney transplantation, was analyzed, with RNAseq data from pre-transplantation kidney biopsies compared to those from post-transplantation biopsies. All relevant plots and analyses were generated using RStudio 2024.12.1+563 software, and only genes with statistically significant differences between groups were included in subsequent analyses.

### Statistical analysis

All statistical analyses were conducted with GraphPad Prism 10 (GraphPad Software, USA). Specific details regarding the statistical tests are provided in the figure legends. *p-*value <0.05 was considered statistically significant.

## Reporting summary

Further information on research design is available in the Nature Portfolio Reporting Summary linked to this article.

## Data availability

All data and information pertinent to this study are included in the main article and Supplementary Material. Additional information and datasets analyzed in this study can be requested from the corresponding author. Source data are provided with this paper.

## Supplementary materials

**Supplementary Table 1.**
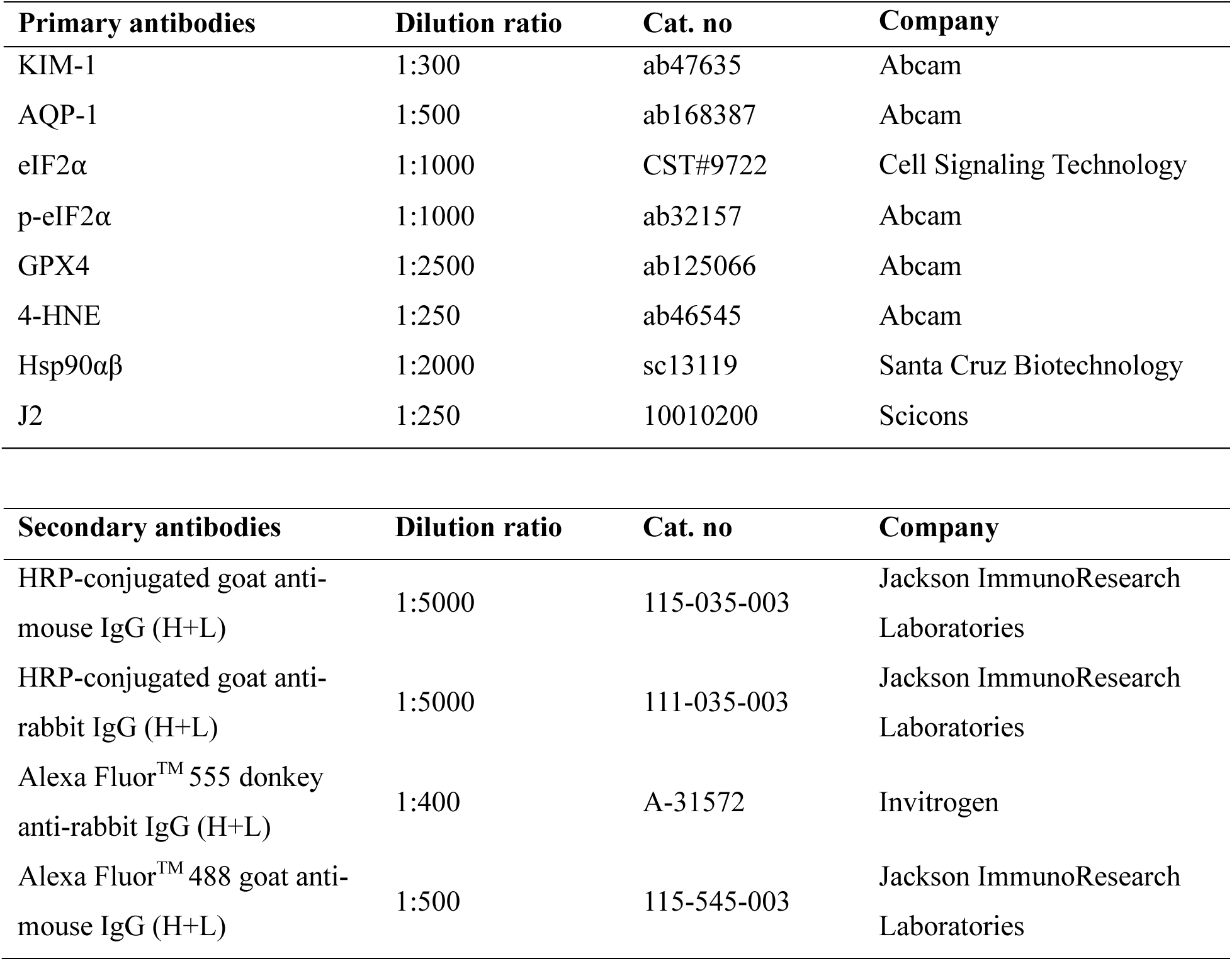
Antibody list utilized in the study.

| Primary antibodies | Dilution ratio | Cat. no | Company |
| --- | --- | --- | --- |
| KIM-1 | 1:300 | ab47635 | Abcam |
| AQP-1 | 1:500 | ab168387 | Abcam |
| eIF2α | 1:1000 | CST#9722 | Cell Signaling Technology |
| p-eIF2α | 1:1000 | ab32157 | Abcam |
| GPX4 | 1:2500 | ab125066 | Abcam |
| 4-HNE | 1:250 | ab46545 | Abcam |
| Hsp90$\alpha\beta$ | 1:2000 | sc13119 | Santa Cruz Biotechnology |
| J2 | 1:250 | 10010200 | Scicons |

| Secondary antibodies | Dilution ratio | Cat. no | Company |
| --- | --- | --- | --- |
| HRP-conjugated goat anti-mouse IgG (H+L) | 1:5000 | 115-035-003 | Jackson ImmunoResearch Laboratories |
| HRP-conjugated goat anti-rabbit IgG (H+L) | 1:5000 | 111-035-003 | Jackson ImmunoResearch Laboratories |
| Alexa Fluor<sup>TM</sup> 555 donkey anti-rabbit IgG (H+L) | 1:400 | A-31572 | Invitrogen |
| Alexa Fluor<sup>TM</sup> 488 goat anti-mouse IgG (H+L) | 1:500 | 115-545-003 | Jackson ImmunoResearch Laboratories |

**Supplementary Table 2.**
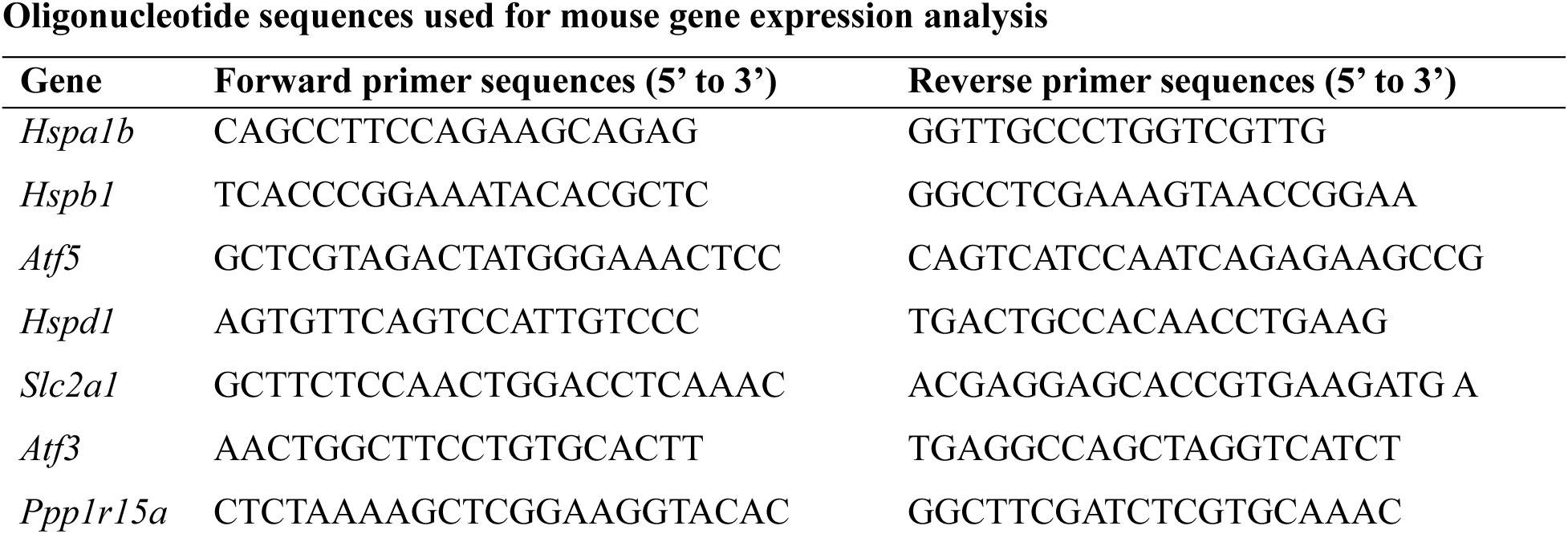
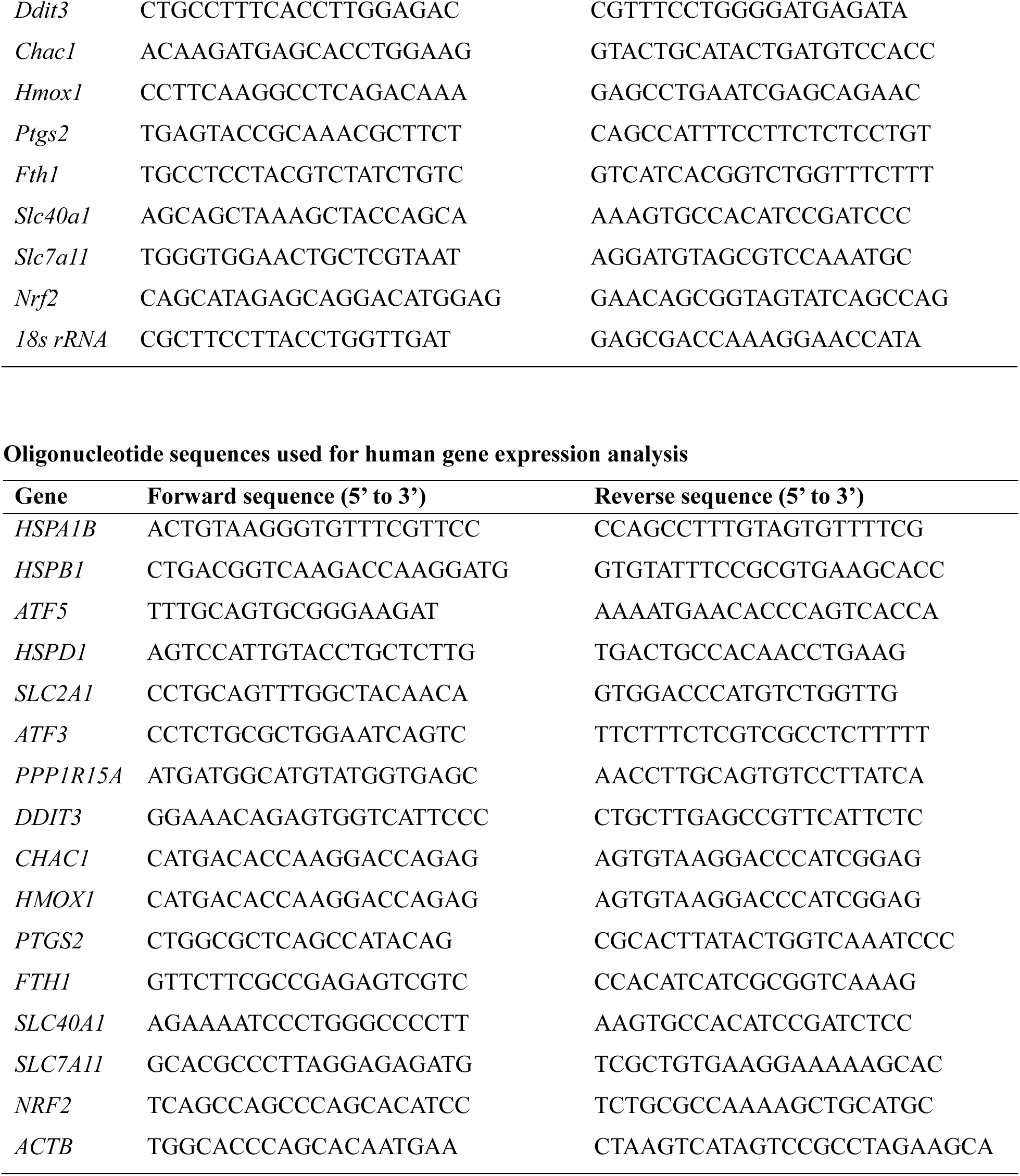
Primer list employed in the study Oligonucleotide sequences used for mouse gene expression analysis.

**Supplementary Table 3.**
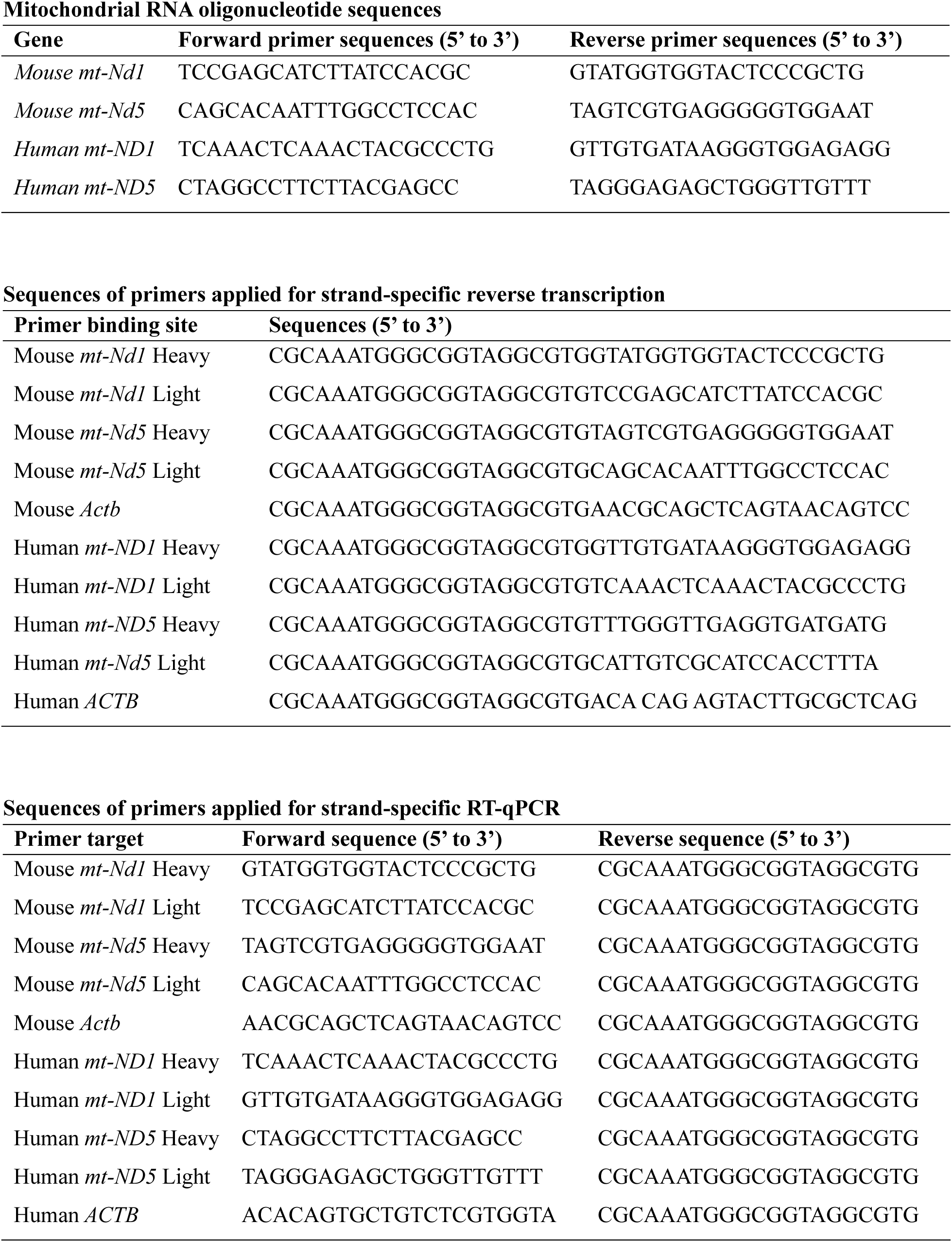
Primer list utilized for mitochondrial dsRNA detection analysis.

**Supplementary Table 4.**
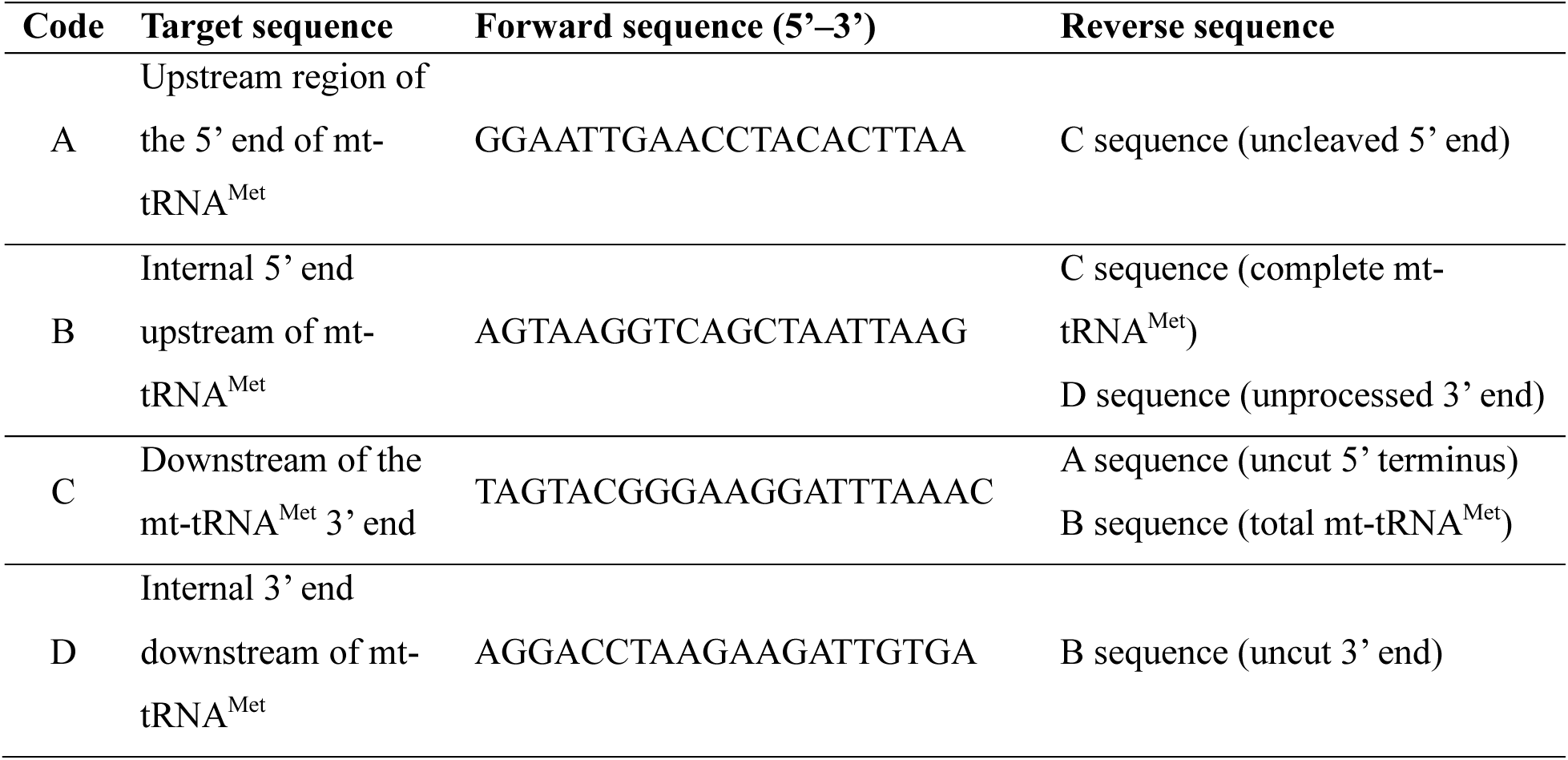
Primer list used for mitochondrial RNA processing analysis.

