## Supplementary for "Mitochondrial Double-Stranded RNA Triggers Ferroptosis via PKR–ISR Signaling in Renal Ischemia–Reperfusion Injury"

### Extended Figures

**Extended Data Fig. 1**

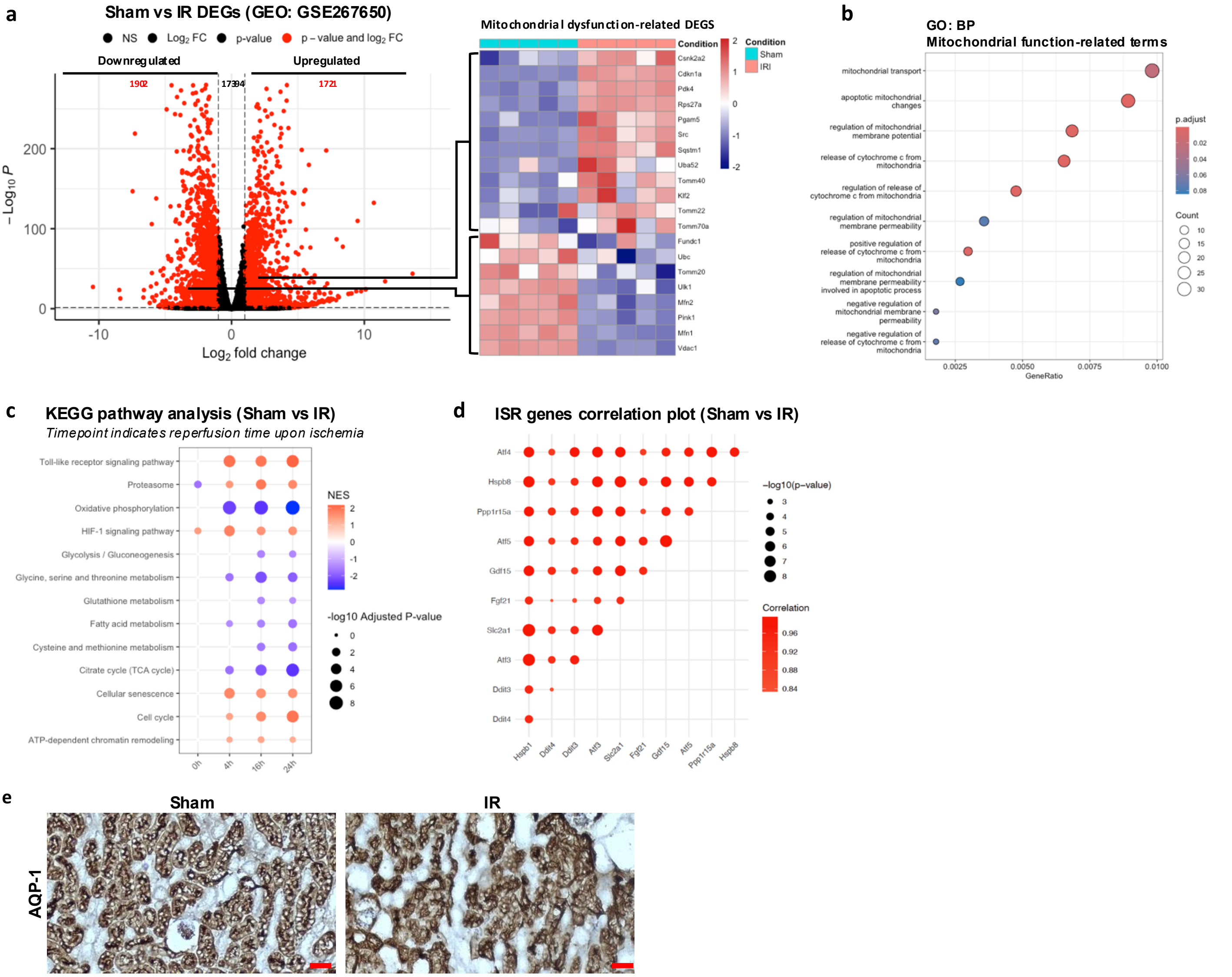

**a,** Differentially expressed genes (DEGs) upon renal ischemia–reperfusion (IR)-treated mouse compared to sham-treated from GSE267650 dataset are shown. Genes associated with mitochondrial dysfunction were identified by comparing sham- and IR-treated mouse, highlighting mitochondrial-specific transcriptional responses to kidney IR injury. **b,** Gene ontology (GO) analysis on biological process revealed significant pathway alterations from mitochondrial function categories. **c,** Enrichment analysis on mitochondrial function-related pathways showed alterations upon gradient reperfusion time post-ischemia in kidney IR-induced mouse. **d,** Pairwise correlation analysis of ISR activated genes during IR, highlighting coordinated transcriptional responses and co-regulatory relationships within the mitochondrial stress network. **e,** Kidney proximal tubule morphological structure between sham and IR-induced mouse group represented by AQP-1. Scale bar = 30 µm.

**Extended Data Fig. 2**

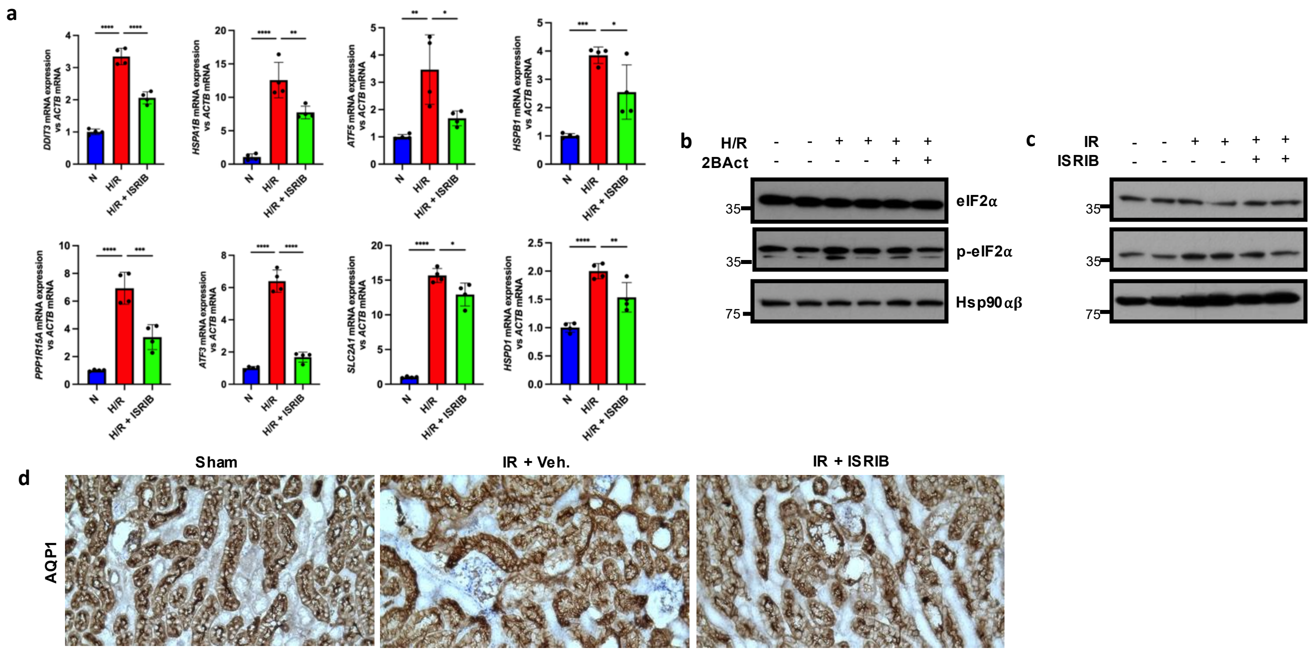


**a,** Gene expression level of ISR-related genes on HK-2 cells after 6 hours hypoxia and 1 hour reperfusion using eIF2B selective inhibitor ISRIB. **b,** Expression level of p-eIF2α upon 2BAct co-treatment 1 hour prior to H/R procedure and during reperfusion period on HK-2 cells. **c,** Expression level of p-eIF2α upon ISRIB treatment 24 hour and 1 hour prior to IR procedure on IR-induced kidney mouse model. **d,** Kidney proximal tubule morphological structure between sham-, IR-, and ISRIB-treated IR kidney injury treatment showed a preserved structure by ISRIB administered, represented by AQP-1 staining.

**Extended Data Fig. 4**

**a**

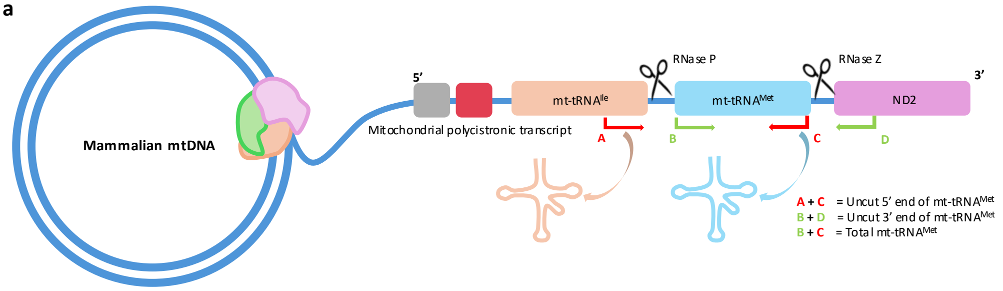


**a,** Experimental diagram for detection of failed mitochondrial RNA processing using primer-specific RT-qPCR analysis. In order for the mitochondrial mRNA to be properly translated from the polycistronic structure, the tRNAs bounding between the mRNAs need to be cut through RNA processing machinery. To achieve this, RNA processing machinery of mitochondrial tRNA$^{Met}$ was targeted to determine the uncut portion from the polycistronic structure. Primers were designed as follow; Uncut 5' end of mt-tRNA$^{Met}$ (A,C): targeting a specific 3' end sequence from the mt-tRNA$^{Ile}$ and a specific 3' end sequence from mt-tRNA$^{Met}$; Uncut 3' end of mt-tRNA$^{Met}$ (B,D): targeting a specific 5' end sequence from mt-tRNA$^{Met}$ and a specific 5' end sequence from mt-ND2. The unprocessed region of mt-tRNA$^{Met}$ were normalized to toal mt-tRNA$^{Met}$ present in the mitochondrial transcript, represented by primer targeting the 5' end and 3' end sequence of mt-tRNA$^{Met}$ (B,C).

**Extended Data Fig. 5**

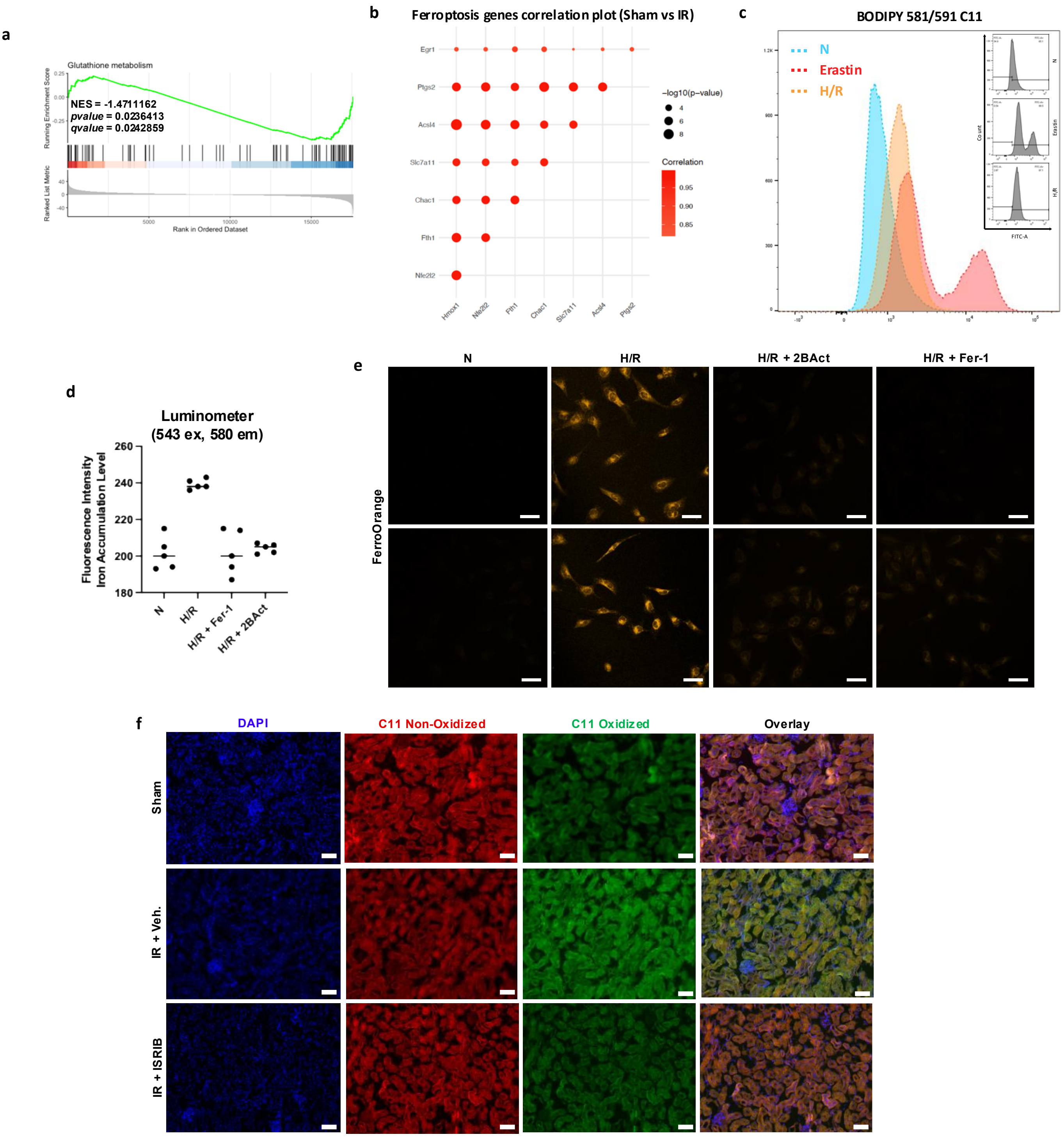


**a,** Gene set enrichment analysis (GSEA) from RNA-Seq dataset (GSE2676750) showing the downregulation of glutathione metabolism as one of the ferroptosis hallmark on IR-induced kidney injury compared to sham-treated mouse, supporting ferroptosis activation. **b,** Pairwise correlation analysis of ferroptosis activated genes during IR, highlighting coordinated transcriptional responses and co-regulatory relationships within the ferroptosis signaling network. **c,** Flow cytometry result showing elevated level of C11 oxidized form of BODIPY 581/591 C11 upon Erastin as the positive marker of ferroptosis inducer and H/R-induced HK-2 cells, representing ferroptosis activation during H/R procedure. **d,** Luminometer reading of FerroOrange to represent the iron accumulation during H/R followed by co-treatment with Fer-1 as ferroptosis inhibitor and 2BAct as ISR inhibitor, showing less accumulated iron upon the treatment on HK-2 cells. **e,** Confocal microscopy visualization of FerroOrange staining showing the accumulation of iron on H/R-induced HK-2 cells and diminished iron level upon co-treatment with 2BAct and Fer-1. Scale bar = 50 µm. **f,** BODIPY 581/591 C11 staining on fresh kidney section upon IR-induced kidney injury and ISRIB co-treatment on IR-induced mouse compared to sham-treated mouse. Scale bar = 90 µm.
